# Embryonic stem cells commit to differentiation by symmetric divisions following a variable lag period

**DOI:** 10.1101/2020.06.17.157578

**Authors:** Stanley E Strawbridge, Guy B Blanchard, Austin Smith, Hillel Kugler, Graziano Martello

## Abstract

Mouse embryonic stem (ES) cells are derived from the epiblast of the preimplantation embryo and retain the capacity to give rise to all embryo lineages. ES cells can be released into differentiation from a near-homogeneous maintenance condition. Exit from the ES cell state can be accurately monitored using the Rex1-GFPd2 transgenic reporter, providing a powerful system for examining a mammalian cell fate transition. Here, we performed live-cell imaging and tracking of ES cells during entry into differentiation for 48 hours in defined conditions. We observed a greater cell surface area and a modest shortening of the cell cycle prior to exit and subsequently a reduction in cell size and increase in motility. We did not see any instance of cells regaining ES cell identity, consistent with unidirectional developmental progression. Transition occurred asynchronously across the population but genealogical tracking revealed a high correlation in cell-cycle length and Rex1-GFPd2 expression between daughter cells. A population dynamics model was consistent with symmetric divisions during exit from naive pluripotency. Collapse of ES cell identity occurred acutely in individual cells but after a variable delay. The variation in lag period can extend up to three generations, creating marked population asynchrony.

## INTRODUCTION

Embryonic stem (ES) cells are derived from the pre-implantation mouse epiblast (Evans and Kaufman, 1981; Martin, 1981) and retain a high level of similarity to their in vivo counterpart (Boroviak et al., 2014). The defining features of ES cells are self-renewal and naïve pluripotency (Jaenisch and Young, 2008; Martello and Smith, 2014; Nichols and Smith, 2012; Smith, 2001). Naïve pluripotency is the latent potential to give rise to all tissues in the adult body, including germ cells (Nichols and Smith, 2009; Smith, 2001) and is governed by a highly recursive transcription factor network (Dunn et al., 2014; Jaenisch and Young, 2008; Niwa, 2018; Young, 2011). Maintenance of the naïve transcription factor network can be achieved by culturing ES cells in the presence of the cytokine leukaemia inhibitory factor (LIF) (Smith et al., 1988; Williams et al., 1988) plus serum, or using two small molecule inhibitors (2i), CHIR99021, a glycogen synthase kinase-3 (GSK3) inhibitor, and PD0325901, a MEK inhibitor that blocks Erk1/2 signalling (Wray et al., 2010; Ying et al., 2008).

Self-renewal is the ability of a cell to give rise to at least one daughter cell of equal potency and is a defining feature of all stem cells (Smith, 2006). In general, self-renewal divisions can be performed either symmetrically, where both daughter cells retain the potency of the mother cell, or asymmetrically, where one daughter cell retains the potency of the mother cell and the other cell takes on a new identity (Morrison and Kimble, 2006). In the case of differentiation, which may also occur symmetrically or asymmetrically, one or both daughter cells will exhibit altered identity.

Single-cell imaging and tracking have been performed to examine molecular inheritance patterns of ES cells culture in serum/LIF conditions (Filipczyk et al., 2015, 2015) and division symmetry of ES cells exiting from LIF+BMP4 cultures (Jasnos et al., 2013; Nakamura et al., 2018). In these conditions, ES cells are heterogeneous, displaying mosaic and dynamic expression of various transcription factors (Chambers et al., 2007; Filipczyk et al., 2015; Hayashi et al., 2008; Toyooka et al., 2008). Notably, ES cells in serum and LIF switch between Nanog high and low states (Herberg et al., 2016). Comparisons of sister cells show a greater similarity during self-renewal than during differentiation in both Nanog-GFP (green fluorescent protein) expression levels (Nakamura et al., 2018) and in single-cell qPCR of microdissected sister cells (Jasnos et al., 2013). However, in both instances, heterogeneity of the starting cell population creates an increased potential for neighbour effects. To determine the extent to which behaviours are driven by intrinsic or extrinsic cues it is advantageous to minimise variability in the starting cell population and to use defined culture conditions.

Following exchange from 2i into N2B27 medium without serum or growth factors, ES cells remain able to re-enter self-renewal for a period before they permanently down-regulate the naïve transcription factor network (Fig. 1A) (Betschinger et al., 2013; Kalkan et al., 2017; Leeb et al., 2014; Martello and Smith, 2014). In this phase of reversibility, reapplication of 2i is sufficient for recovery of ES cell identity and self-renewal. Cells are deemed to have exited the ES cell state at the point when self-renewal potential is lost (Betschinger et al., 2013; Kalkan and Smith, 2014). Although ES cell populations in 2i are substantially homogeneous and described as residing in a ground state, exit is observed to occur asynchronously over a period of around 48hrs (Betschinger et al., 2013; Kalkan et al., 2017; Mulas et al., 2017). Asynchronicity in early development has been documented in vivo, during acquisition of the naïve state (Saiz et al., 2016) and dissolution of the naïve network (Acampora et al., 2016; Malaguti et al., 2019; Neagu et al., 2020). However, it is unknown how asynchronous behaviour arises and whether it involves asymmetric cell divisions.

**Figure 1.**
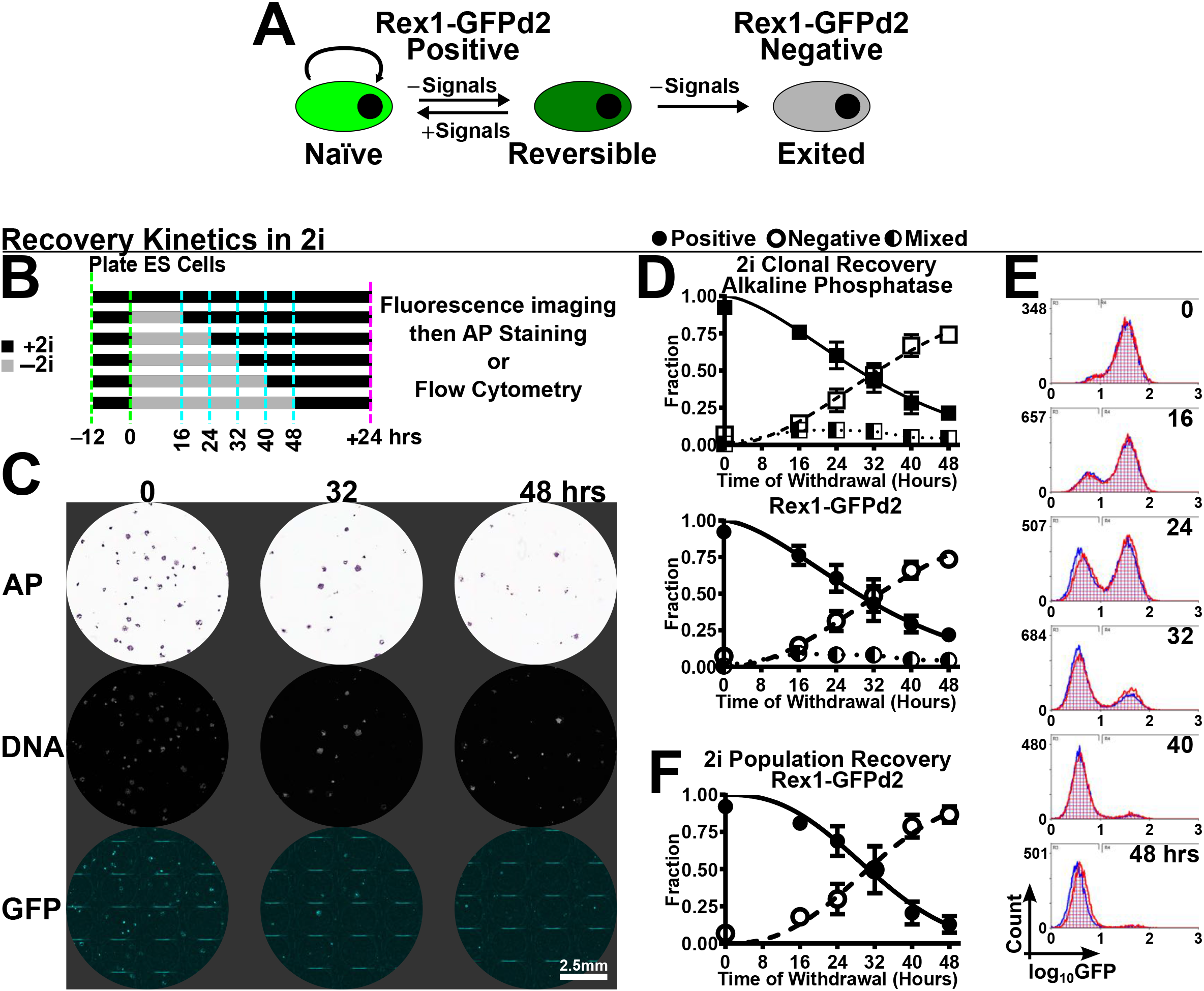
**A.** ES cells can be maintained in the naïve pluripotent state by exogenous signals. Upon signals withdrawal, ES cells dwell in a reversible phase, where signals can be reapplied and cells return to the naive state. Prolonged signals withdrawal results in irreversible exit and reapplication of signals has no effect. Rex1-GFPd2 allows monitoring in near real time the exit dynamics. **B.** Experimental strategy used to measure the dynamics of exit from naive pluripotency in cells cultured in 2i. **C.** Representative images of recovery kinetics assays at clonal density after the indicated hours of 2i withdrawal. Colonies were scored both by alkaline phosphatase staining (AP, top row) and by Rex1-GFP1d2 fluorescence (bottom row). Nuclei staining by Hoechst33342 (middle row) allowed the identification of GFP negative colonies. **D.** The fraction of each colony type is shown for alkaline phosphatase (squares in top panel) and Rex1-GFPd2 (circles in bottom panel). Each time-point is the mean±SEM of 8 biological replicates, from 4 independent experiments, where filled markers show positive colonies, empty markers show negative colonies, and half-filled markers show mixed colonies. The Weibull distribution (*CDF*) was fit to both the positive fraction (solid line) and negative fraction (dashed line) of colonies. The dotted line connects the mixed fraction of colonies. **E.** Flow cytometry profiles from the population recovery kinetics. Two biological replicates from a representative experiment are shown in red and blue. x-axis: log_10_GFP. y-axis: count. Each profile consists of at least 10,000 cells. **F.** Quantification of the flow cytometry profiles from population recover kinetics showing the fraction of Rex1-GFPd2 positive (filled markers) and negative (empty markers) cells. Each time-point is the mean±SEM of 8 biological replicates from 4 independent experiments. The Weibull distribution was fit to both the positive fraction (solid line) and negative fraction (dashed line) of cells.

Rex1 (gene name *Zfp42)* is a transcription factor whose expression is tightly linked to the naïve pluripotency network (Wray et al., 2010). Rex1 linked fluorescent reporters have been widely used as tools that demarcate naïve pluripotency (Guo et al., 2010; Herberg et al., 2016; Nakai-Futatsugi and Niwa, 2016; Toyooka et al., 2008; Wray et al., 2010). Use of destabilised GFP with a 2 hour half-life (Rex1-GFPd2) allows for accurate estimation of exit from naïve pluripotency (Betschinger et al., 2013; Leeb et al., 2014; Mulas et al., 2017, 2017). Here we exploited this system to track exit from naïve pluripotency in defined conditions by time-lapse imaging. During the exit process, we observed a shortening of the cell-cycle, larger cell sizes, and increased motility. As expected, the time of exit was asynchronous at the population level, however, high levels of synchronicity were exhibited by cells within lineages, but not between lineages. In fact, genealogical analyses, supported by a population dynamics model, show that ES cells exit naïve pluripotency using symmetric divisions. Finally, we did not observe spontaneous reactivation of the Rex1-GFPd2 reporter upon deactivation, rather, once cells begin to exit naïve pluripotency they do so at a fixed and rapid rate.

## RESULTS

### *Exit from* naïve *pluripotency remains asynchronous at clonal cell density*

It has previously been noted that ES cell progression from the 2i ground state is asynchronous at the population level (Kalkan et al., 2017; Mulas et al., 2017). Cell-cell interactions could contribute to asynchronous behaviours or otherwise influence transition dynamics. We therefore first assessed exit dynamics at clonal density. We used loss of self-renewal ability upon restoration to 2i as a functional measure of departure from the ES cell state (Betschinger et al., 2013). Prior culture in LIF has been shown to delay exit (Dunn et al., 2014). Therefore, we initiated all experiments from ES cells in 2i alone (Dunn et al., 2014; Mulas et al., 2017).

Rex1-GFPd2 ES cells were plated in 2i on laminin at either 500 cells/10 cm^2^, termed “clonal density”, or at 10-fold higher density (5000 cells/10 cm^2^) (Fig. 1B). After 12 hours, medium was replaced with N2B27 alone to release cells from the ground state. At different timepoints post-withdrawal 2i was restored. Cultures were maintained for 84 hours in total before assay. For the clonal density assay, individual colonies were imaged for nuclei (Hoechst), GFP, and alkaline phosphatase (AP) staining, while flow cytometry was applied to quantify GFP expressing cells for the higher density plates (Fig. 1B).

Colony types were scored as positive, mixed, or negative based on AP staining and GFP fluorescence (Fig. 1C and Fig. S1A). A small fraction of negative colonies were apparent after 2i withdrawal for 16h and this number increased dramatically at 24 and 36 hours (Fig. 1C,D). By 48 hours, the majority of colonies (79±3%) were negative for both AP and GFP. Exit from naïve pluripotency can be thought of as collapse, or failure, of a transcription factor network. The Weibull distribution has been widely used to model the failure of processes (Weibull, 1951). We, therefore, fitted exit timings to a 2-parameter Weibull cumulative distribution function (*CDF*) (Fig. 1D, see Methods). The shape parameter (γ) of the Weibull distribution allows us to infer how the process evolves in time. The mean(±SD) time-of-exit was approximately 32(±20) hours (Supplemental Table 1) with both AP and GFP markers showing the same kinetics (p-value = 1 for mean and p-value = 1 for SD; t-Test with Bonferroni post hoc correction, Supplemental Table 2). The shape parameter (γ =1.61) was greater than one, implying that collapse of the pluripotency network increases over time.

For the higher density cultures (Fig. 1E), we resolved GFP positive and negative populations by flow cytometry (Fig. S1B). The time course of decline in the GFP positive fraction was similar to that for loss of ES cell colony formation at clonal density, with 87±6% of the population GFP negative by 48h (Fig. 1F). Fitting the Weibull distribution gave a mean(±SD) time-of-exit of 32(±13) hours with a shape parameter greater than one (γ =2.5). The two parameters were not significantly different between the two cell densities (Supplemental Table 2).

These analyses showed that exit from pluripotency occurs over 48 hours with individual cells exiting at different timepoints, consistent with previous observations (Betschinger et al., 2013; Kalkan et al., 2017; Leeb et al., 2014; Mulas et al., 2017). Importantly these dynamics were maintained at clonal density, suggesting intrinsic control.

### Single-cell tracking of ES cell exit from naïve pluripotency

Preservation of transition dynamics at clonal density allowed us to apply long-term single-cell imaging and tracking. To facilitate cell segmentation, we introduced a plasma membrane fluorescent reporter into Rex1-GFPd2 cells. For this purpose, we created a fusion between the palmitoylation sequence of Gap43 and the N-terminus of mCherry (Fig. 2A), to produce a reporter that is localised to the interior of the plasma membrane via two lipid tails (Goedhart et al., 2012). Following transfection and selection we isolated a Rex1-GFPd2/Gap43-mCherry clone with sufficiently intense signal to demarcate cell-cell boundaries in ES cells (Fig. 2B). This clone displayed similar exit kinetics to parental Rex1-GFPd2 cells assayed by flow cytometry (p-value = 0.6892, two-way ANOVA) (Fig. 2C).

**Figure 2.**
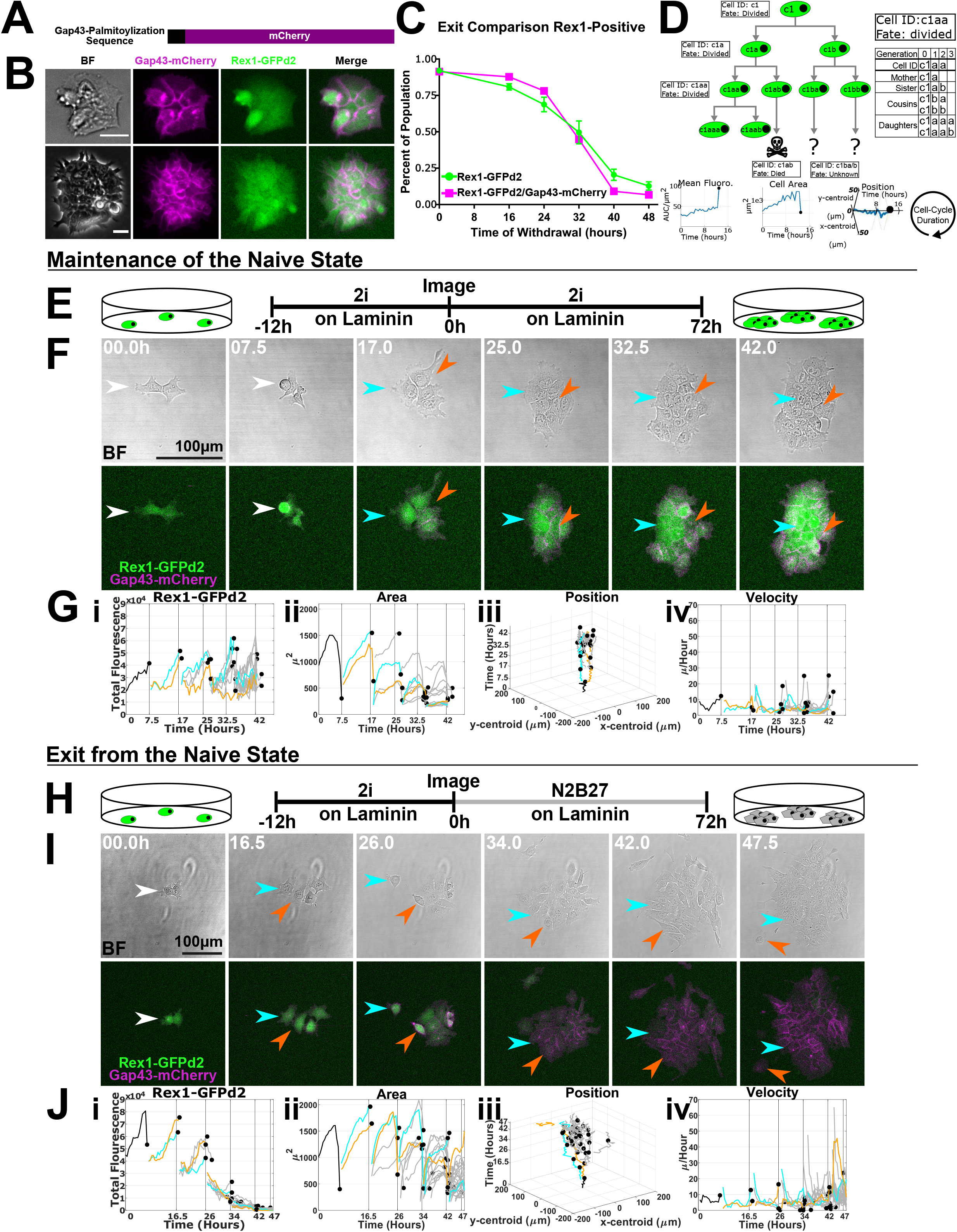
**A.** mCherry is N-terminally tagged with the palmitoylation sequence of Gap43. Cysteines 3 and 4 are the residues of Gap43 which are modified with palmitate which produces a mCherry that will be attached to the interior of the plasma membrane. **B.** Images of the dual reporter Rex1-GFPd2/Gap43-mCherry ES cells in 2i on laminin. The two rows show two different colonies each with brightfield (BF), membrane localised Gap43-mCherry and the Rex1-GFPd2 reporter (scale bars=25 um). **C.** Validation for Rex1-GFPd2/Gap43-mCherry ES Cells by population level recovery kinetics. The parental Rex1-GFPd2 ES cell line (green line, n=8) and its subclone Rex1-GFPd2/Gap43-mCherry ES cell line (magenta line, n=2) shows no difference in kinetics (p-value = 0.6892, two-way ANOVA). Markers show mean with error bars showing SEM. **D.** Schematic of the information generated by the single-cell tracking for lineages and cells. **E.** Experimental strategy to perform long-term single-cell imaging of ES cells maintained in the naive state. **F.** Representative time-course showing BF (top row) and Rex1-GFPd2/Gap43-mCherry (bottom row) with arrows indicating cells from a single branch of the lineage tree (See panel G). **G.** Quantified metrics from the lineage in panel F with colour corresponding to arrows and vertical black lines corresponding to time points. **H.** Experimental strategy to perform long-term single-cell imaging of ES cells exiting the naive state. **I.** Representative time-course showing BF (top row) and Rex1-GFPd2/Gap43-mCherry (bottom row) with arrows indicating cells from a single branch of the lineage tree (See panel J). **J.** Quantified metrics from the lineage in panel I with colour corresponding to arrows and vertical black lines corresponding to time points.

Rex1-GFPd2/Gap43-mCherry cells were plated on laminin at clonal density in 2i. Twelve hours after plating, the medium was replaced with fresh 2i (Fig. 2E-G and Fig. S2B) or with N2B27 (Fig. 2H-J). E14Tg2a ES cells transfected with Gap43-mCherry were used as a negative control for the GFP signal (Fig. S2A). Confocal z-stacks of multiple positions were collected for brightfield, GFP, and mCherry every 30 minutes for a total of 72 hours. The GFP z-stack images were summed and the cells in the time-course stacks were manually tracked and cell shapes recorded (England et al., 2006). Cell tracks and shapes were fed into tracks analysis software oTracks (Blanchard et al., 2009). An additional analysis module was written to quantify the resulting manual tracks for position, area, fluorescence, and genealogy (Fig. 2D). Terminal fates were assigned to each cell as either ‘divided’, ‘died’, or ‘unknown’. The ‘unknown’ cell fate was prescribed when a cell moved out of frame or could no longer be identified reliably due to excessive crowding.

In 2i, cells maintained robust GFP signal throughout the entire time-course (Fig. 2E-G), indicating that the imaging conditions did not reduce cell viability and did not cause photobleaching of GFP (Fig. S2B). We observed that GFP fluorescence behaved seasonally over the cell-cycle. After division, the signal was initially low and then increased gradually until the next division. This observation was consistent with previous long-term imaging studies (Filipczyk et al., 2015; Herberg et al., 2016; Nakai-Futatsugi and Niwa, 2016). Under exit conditions (Fig. 2H-J), we observed the expected reduction in GFP signal from 16 hours onwards. Figure S2C reports the sample size for the two biological replicates for the analysis of the long-term single-cell tracking data. In total, 2,624 individual cells from two independent experiments were tracked up to the fifth generation with confidence.

### Exit from naïve pluripotency is associated with a shorter cell-cycle, transient cell spreading, and increased motility

From long-term single-cell imaging data we were able to measure the cell-cycle length directly, as the time between two cell divisions. We found that the cell cycle duration in 2i (10.20±2.14 hours) was longer than for cells exiting in N2B27 (9.26±2.03 hours) (p-value = 2.771×10^-9^: Rank Sum) (Fig. 3A). The cell-cycle duration of ES cells in 2i remains constant over generations (p-value = 0.2269: Kruskal-Wallis) in contrast to exiting cells in N2B27 that show a progressively shortening cell-cycle length (p-value =1.29×10^-47^) (Fig. 3B). Similar results were obtained from the analysis of static images of ES cell colony growth from single cells after 2i withdrawal (Fig. S3A-F), further indicating that exit from naïve pluripotency is associated with cell-cycle shortening.

**Figure 3.**
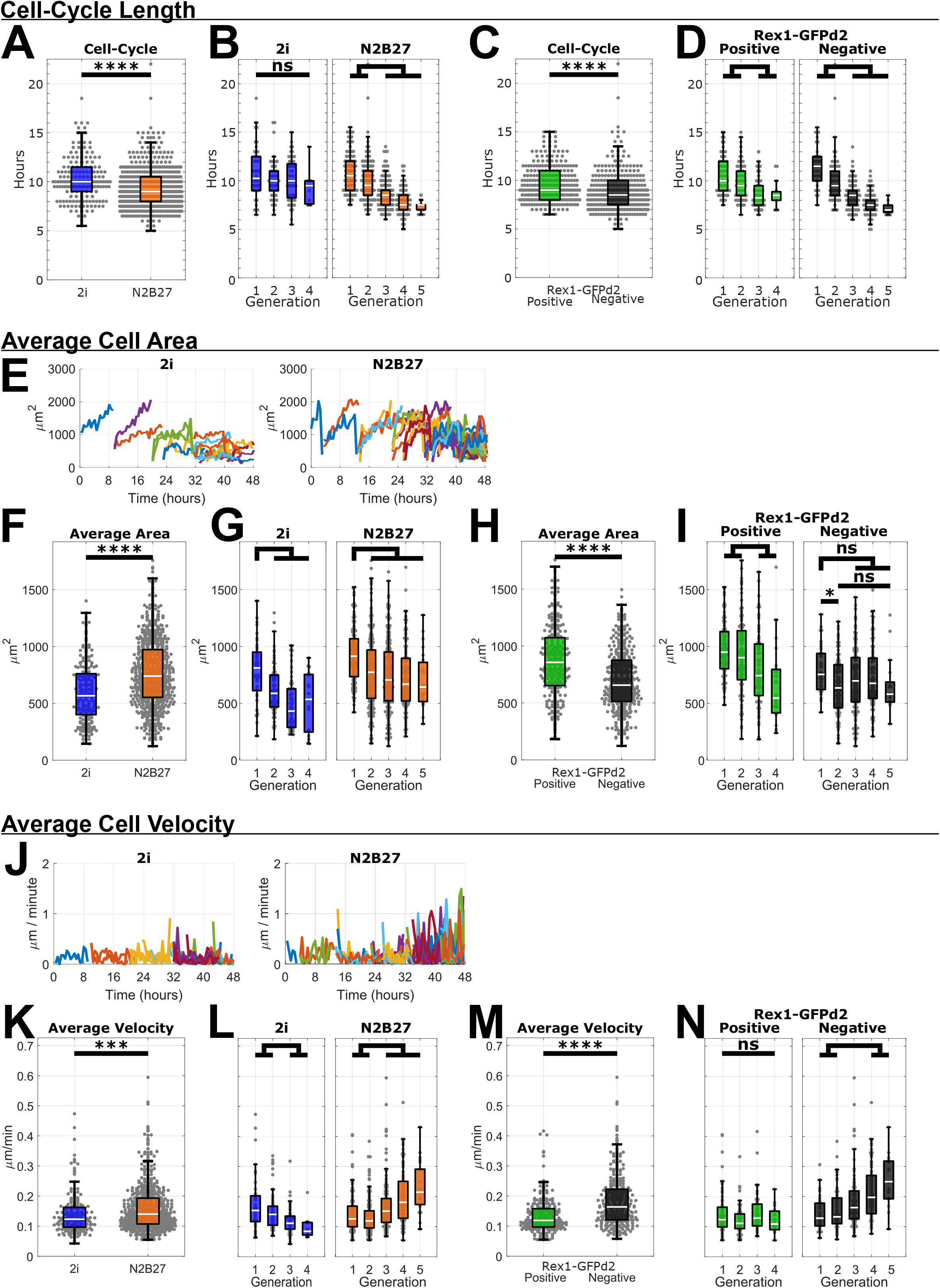
**A-D, F-I, K-N.** Box plots superimposed with swarm plots where each point shows measurements for a single cell. All analyses shown are from Experiment 1 (see Figure 3K) and similar results were obtained from Experiment 2. **A.** The cell-cycle length of ES cells in 2i is 10.20±2.14 hours (N_cells_= 196). This is one hour longer than cells in N2B27, which have a cell-cycle length of 9.26±2.03 hours (N_cells_= 602) (p-value = 2.771×10^-9^, two-tailed unpaired Wilcoxon Rank Sum). **B.** ES cells in 2i have constant cell-cycle over subsequent generations (p-value = 0.2269, Kruskal-Wallis Test with Bonferroni post-hoc correction), where cells in N2B27 show a trend of a decreasing cell-cycle length (p-value =1.291×10^-47^, Kruskal-Wallis Test). **C.** Rex1-GFPd2 positive cells have longer cell-cycle, of 9.68±1.87 hours (N_cells_= 275) than Rex1-GFPd2 negative cells, with a cell-cycle of 8.91±2.10 hours (N_cells_= 327). (p-value = 1.108×10^-8^, two-tailed unpaired Wilcoxon Rank Sum). **D.** There is an overall shortening of cell-cycle over subsequent generations for both Rex1-GFPd2 positive cells (p-value =5.209×10^-11^, Kruskal-Wallis Test with Bonferroni post-hoc correction) and Rex1-GFPd2 negative cells (N2B27, p-value =1.078×10^-29^, Kruskal-Wallis Test with Bonferroni post-hoc correction). **E.** Representative lineages for 2i and N2B27 where each line represents a cell. **F.** ES cells in 2i have a smaller cell area, of 596±257 μm^2^ (N_cells_=207), than those exiting in N2B27, which have a cell area of 768±303 μm^2^ (N_cells_=725) (p-value = 4.071×10^-13^, two-tailed unpaired Wilcoxon Rank Sum). **G.** The area a cell occupies decreases over subsequent generations for ES cells in 2i (p-value = 9.600×10^-8^, Kruskal-Wallis Test with Bonferroni post-hoc correction) and cells exiting in N2B27 (p-value =5.693×10^-13^, Kruskal-Wallis Test). **H.** Rex1-GFPd2 positive cells have a larger cell areas, of 873±318 μm^2^ (N_cells_=304), than Rex1-GFPd2 negative cells, with an area of 693±268 μm^2^ (N_cells_=421) (p-value = 2.609×10^-14^, two-tailed unpaired Wilcoxon Rank Sum). **I.** The area a cell occupies decreases over subsequent generations for Rex1-GFPd2 positive cells (p-value = 9.600×10-8, Kruskal-Wallis Test with Bonferroni post-hoc correction). However, for cells exiting in N2B27, a different cell area is only seen between the first and second generation (generation 1vs2: p-value =0.0436; all other comparisons: p-values>0.05, Kruskal-Wallis Test with Bonferroni post-hoc correction). **J.** Representative lineages for 2i and N2B27 where each line represents a cell. **K.** Average cell velocity of ES cells in 2i, 0.14±0.07 μm/min (N_cells_=196), is slower than those exiting in N2B27, 0.16±0.08 μm/min (N_cells_=602) (p-value = 1.391×10^-4^, two-tailed unpaired Wilcoxon Rank Sum). **L.** ES cells in 2i exhibit a decrease in average velocity (p-value = 1.049×10^-®^ Kruskal-Wallis Test with Bonferroni post-hoc correction), where cells exiting in N2B27 (p-value =2.379×10^-15^, Kruskal-Wallis Test) became more motile. **M.** Rex1-GFPd2 positive are also less motile (0.13±0.06 μm/min, N_cells_=275) than the Rex1-GFPd2 negative cells (0.18±0.08 μm/min, N_cells_=327) (p-value =4.414×10^-1®^ two-tailed unpaired Wilcoxon Rank Sum). **N.** No change is seen in the average velocity of Rex1-GFPd2 positive cells (p-value =0.0559, Kruskal-Wallis Test with Bonferroni post-hoc correction). The Rex1-GFPd2 negative population displays an increase in motility (p-value =7.082×10^-9^, Kruskal-Wallis Test with Bonferroni post-hoc correction).

We examined whether cell-cycle shortening begins immediately when cells are withdrawn from 2i or begins later in the exit process. Cell images were parsed into GFP positive and negative using an empirically determined threshold (Fig. S3G). We found that GFP positive cells in N2B27 had a shorter cell-cycle (9.68±1.87 hours) than cells in 2i, but longer than GFP negative cells (8.91±2.10 hours) (Fig. 3C) (p-value = 1.108×10^-8^). Therefore shortening of the cell-cycle begins prior to exit from naïve pluripotency, as recently proposed (Waisman et al., 2019), and the cell cycle shortens further over subsequent generations (Fig. 3D).

We evaluated change in cell size during transition, measured as the area covered by a cell (Fig. 3E). Determination of the average cell area revealed that ES cells in 2i (596±257 μm^2^) are smaller than exiting cells (768±303 μm^2^, p-value = 4.071×10^-13^) (Fig. 3F). In either condition, cell area decreased over generations as the number of cells in a colony increased (Fig 3G). However, discrimination between GFP positive and negative cells during exit identified that GFP positive cells exhibit a greater surface area (873±318 μm^2^) compared with negative cells (693±268 μm^2^) (p-value = 2.609×10^-14^) or 2i ES cells (Fig. 3H). Thus, cells flatten and spread out prior to exit, and subsequently contract. The decrease in cell area occurred between the first and second generation (p-value =0.0436) after loss of GFP, and thereafter cell area remained relatively constant.

We also observed increased motility from around the time that cells down-regulated GFP (Fig. 3J). The average velocity of ES cells in 2i (0.14±0.07 μm/min) was less than in N2B27 (0.16±0.08 μm/min) (p-value = 1.391×10^-4^) (Fig. 3K). Over generations, ES cells in 2i exhibited a decrease in average velocity (p-value = 1.049×10^-6^), whereas cells in N2B27 (p-value =2.379×10^-15^) became more motile (Fig. 3L). In N2B27 GFP positive cells are less motile (0.13±0.06 μm/min) than GFP negative cells (0.18±0.08 μm/min) (p-value =4.414×10^-16^) (Fig. 3M). Moreover, motility increased over generations in the GFP negative population (p-value =7.082×10^-9^) (Fig. 3N).

In summary exit from naïve pluripotency is preceded by a shortened cell cycle, a larger cell surface area and a constant low level of motility. After exit, the cell-cycle shortens further, cell area reduces then remains constant, and motility increases progressively.

### Related cells exit naïve pluripotency at similar times

To gain insight into the asynchronous nature of the ES cell transition we investigated whether exit dynamics were more similar between lineally related than non-related cells. From the trajectory of the GFP signal for each cell we extracted four parameters: initial value, final value, mean value, and integrated area under the curve (AUC) (Fig. 4A). We then performed genealogical comparisons within generations (Fig. 4B) between sister cells (ss) and first-cousins (cc), and across generations (Fig. 4C) between mothers and daughters (md) and grandmothers and granddaughters (gd). For all comparisons we calculated the Rank Sum p-value, which takes high values when cells are similar, or low values when cells are significantly dissimilar. In the case of final GFP level (Fig. 4D-F) we saw that in 2i all cells are GFP positive (Wray et al., 2010), leading to a high level of similarity between sisters (Rank Sum p-value=0.2193 for ss).-In N2B27, ss and cc (Fig. 4E,F) showed little difference between pairs of cells (Rank Sum p-value=0.7416 and 0.3388, respectively). Significant differences between pairs were seen only for both md (Rank Sum p-value=8.372×10^-22^) and gd (Rank Sum p-value=3.381×10^-58^).

**Figure 4.**
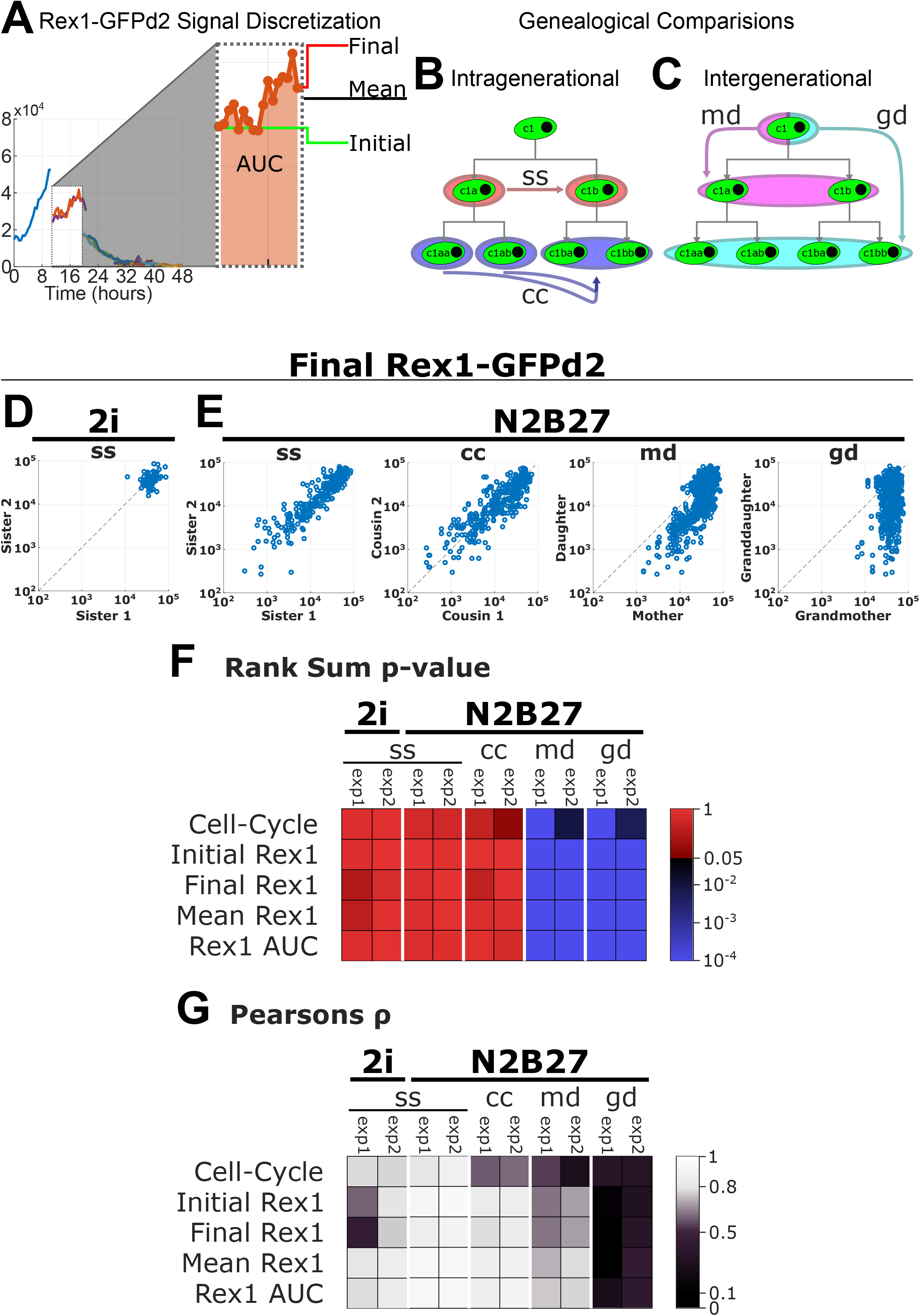
**A.** The Rex1-GFPd2 trajectory for each cell was discretized into the initial, final, mean, and area under the curve (AUC). **B.** Intra-generational comparisons of metrics were performed for sister (ss) and first cousin (cc) pairs. **C.** Inter-generational comparisons of metrics were performed for mother-daughter (md) and grandmother-granddaughter (gd) pairs. **D.** Sister (ss) comparison for final Rex1-GFPd2 level in 2i. n= 82 comparisons shown as dots. Rank Sum p-value=0.2193, Pearson ρ=0.4360, as determined by permutation test. **E.** Genealogical comparisons for Rex1-GFPd2 level in N2B27, comparisons shown as dots. Sister (ss): Rank Sum p-value=0.7416, Pearson ρ=0.8626, n=271. First cousin (cc): Rank Sum p-value=0.3388, Pearson ρ=0.7835, n=386. Mother-daughter (md): Rank Sum p-value=8.372×10^-22^, Pearson ρ=0.6288, n=602. Grandmother-granddaughter (gd): Rank Sum p-value=3.381×10^-58^, Pearson ρ=0.0494, n=478. **F.** Heatmap showing Paired Rank Sum for the difference in value for 5 different parameters between genealogical relations, calculated for two Independent experiments. The colour map is arranged to delineate significance from non-significance, where blue corresponds to a significant difference between relations and where red corresponds to no significance. **G.** Pearson correlation between genealogical relations calculated for 5 parameters on data from two independent experiments. The colour map is arranged to delineate effect size as prescribed by Cohen (2013). Significance from non-significance, where blue corresponds to a significant difference between relations and where red corresponds to no significance. Black is used for ρ<0 showing no effect. Dark purple shows a small effect with 0.1≤ρ<0.5, light purple shows a medium effect with 0.5≤ρ<0.8, and bright greys show a large effect with 0.8≤ρ.

We performed similar analyses for all four GFP metrics and also cell-cycle length (Fig. 4F,G) and both ss and cc in N2B27 showed no significant difference. We also calculated Pearson values, as an independent metric of similarity of parameters between pairs of cells (Fig 4G). In N2B27 ss and cc exhibited a high level of correlation, often higher than ss in 2i. Lower levels of correlation were observed for md and gd relationships.

For all parameters considered, similarity and correlation were very high both in ss and cc in N2B27. Thus, sister cells are born with the same amount of GFP, they maintain a very similar level over their lives, as indicated by the mean and AUC fluorescence, and at the end of a cell-cycle of nearly identical length, they show the same fluorescence intensity. These observations demonstrate that sister ES cells adopt the same fate and follow similar dynamics during exit from naïve pluripotency.

### Population dynamics modelling substantiates a symmetric division mode during exit from naïve pluripotency

The preceding observations imply that exit from naïve pluripotency proceeds via symmetric divisions. In recent years mathematical modelling has been applied extensively to understand patterns of cell division in adult tissue stem cell populations (Clayton et al., 2007; Lopez-Garcia et al., 2010; Simons and Clevers, 2011). We turned to modelling to examine: (1) how the probability of exit changes over time; (2) the proportion of symmetric and asymmetric divisions during exit. We model the exit from naïve pluripotency in a population of GFP positive *P* and negative *N* cells. In our model, cells divide with a rate *λ*, die at a rate that grows logistically with total cell number (specified by two parameters, δ and *K*) (Fig. 5A), and exit the naïve state (Supplemental Model). The net growth rate *γ* = *λ* – *δ* and the carrying capacity *K* = ((*λ* – *δ*)· *K*_0_)/*δ*, where *K*_0_ determines how strongly cell death depends on cell density. We assert that exit is an irreversible process, and we couple it to the divisions of GFP positive cells (Fig. 5B). In the absence of further data, to model the probability of exit, we choose from a set of phenomenological candidate functions *f* (*P, N, t*). Exit divisions are partitioned into symmetric, with probability *ρ*, and asymmetric, with probability 1 – *ρ*. These dynamics yield a system of coupled first order nonlinear ordinary differential equations,

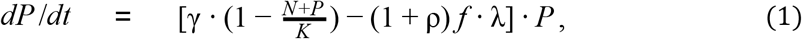

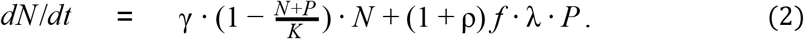

**Figure 5.**
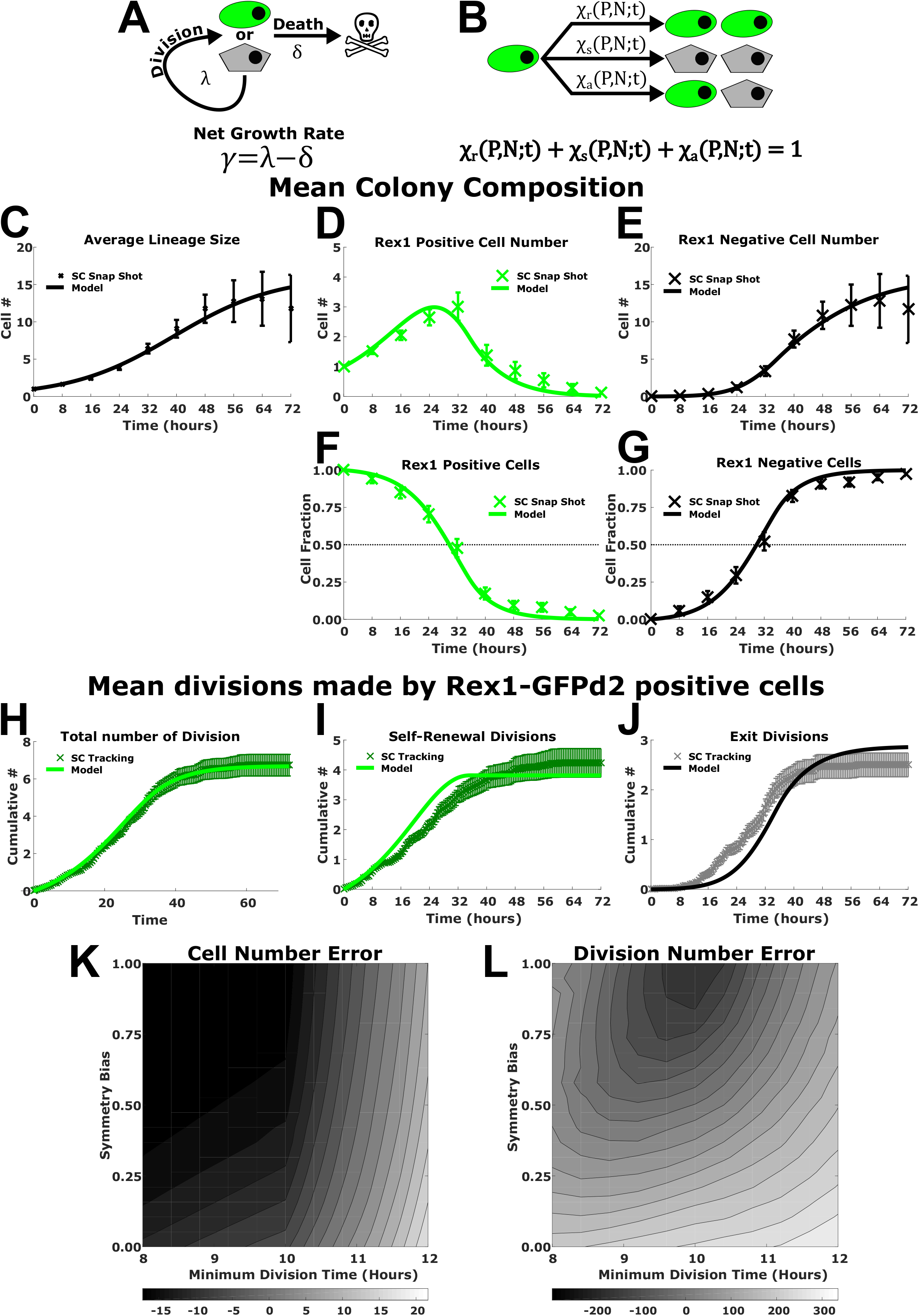
**A.** Our model allows for cells to divide with rate λ and die with rate δ, with a net growth rate of y. Here undifferentiated cells are shown as green ovals and cells that are exited naive pluripotency are shown as grey pentagons. **B.** Divisions for Rex1-GFPd2 cells are partitioned into symmetric self-renewal χ_R_’ symmetric exit χ_S_’ and asymmetric χ_A_. **C-G.** Time-course of colony composition from the single-cell snapshot data used to infer model parameters. The “x” shows the mean and error bars show SEM. Superimposed is the leading-order polynomial model simulated using inferred parameters (solid line). **H-J.** Time-course of divisions made by Rex1-GFPd2 cells from the single-cell tracking data. The “x” shows the mean and error bars show SEM. Superimposed is the leading-order polynomial model simulated using inferred parameters from the single-cell snap-shot data (solid line). **K-L.** Error plots showing the sum squared error between the empirical data and the simulations over the constraints of minimum division time and symmetry bias. Colours indicate sum squared error.

The data used in parameter inference were obtained by counting the number of cells present in each clone at 8 hour intervals for each video, while simultaneously scoring cells as GFP positive or negative. These data (‘x’ in Fig. 5C-G) provide the mean number of cells in a colony at a given time-point and the colony composition. Model parameters were inferred by constrained optimisation, minimising the sum squared error (SSE) between simulations and empirical data.

Using the inferred parameters from Figure 5C-G, we test the model’s predictive power by simulating the total number of divisions (Fig. 5H), self-renewal divisions (Fig. 5I) and exit divisions (Fig. 5J) performed by GFP positive cells and comparing against single-cell tracking data. To investigate what parameter set and test function provides the best agreement between all empirical data (Fig. 5C-J) and the model, we calculated the sum squared error for different paired values of the symmetry bias and cell-cycle length. We found that the optimal parameter set is a cell-cycle length near 10 hours and a **ρ** near 1 (Fig. 5K-L). Thus, the empirical data are consistent with a simple model in which exit divisions occur symmetrically and the probability of exit increases over time.

Finally, we were able to analytically solve the model equations with a time-explicit leading order polynomial (LOP) test function and show that the solution takes the same form as the Weibull distribution (Supplemental Model, Equation 26). Calculating the Weibull parameters from the inferred parameters values of LOP, we arrive at similar values to those determined empirically (Figure 1). The model therefore provides a further support that exit from naïve pluripotency can be viewed as a time-dependent failure process (Supplemental table 1).

### ES cells exit at different timepoints but with similar and rapid half-life decay kinetics

Our model assumes that the probability of exit simply increases over time and does not occur at a specific time after cell division or after a given number of divisions. Indeed, in contrast to cells in 2i, which remain GFP positive over multiple generations (Fig. 6A), cells in N2B27 show down-regulation of GFP at different points in the life-time of the cell, in different generations, and at different times post-2i withdrawal (Fig. 6B, black arrows). Most interestingly, we noted that the GFP signal dropped rapidly immediately before exit, obeying exponential decay (Fig. 6B). We did not observe cells reactivating the GFP reporter after crossing the threshold in 1,957 cells examined up to the fifth generation, confirming the unidirectional nature of the exit process in defined conditions (Kalkan et al., 2017; Smith, 2013, 2017).

**Figure 6.**
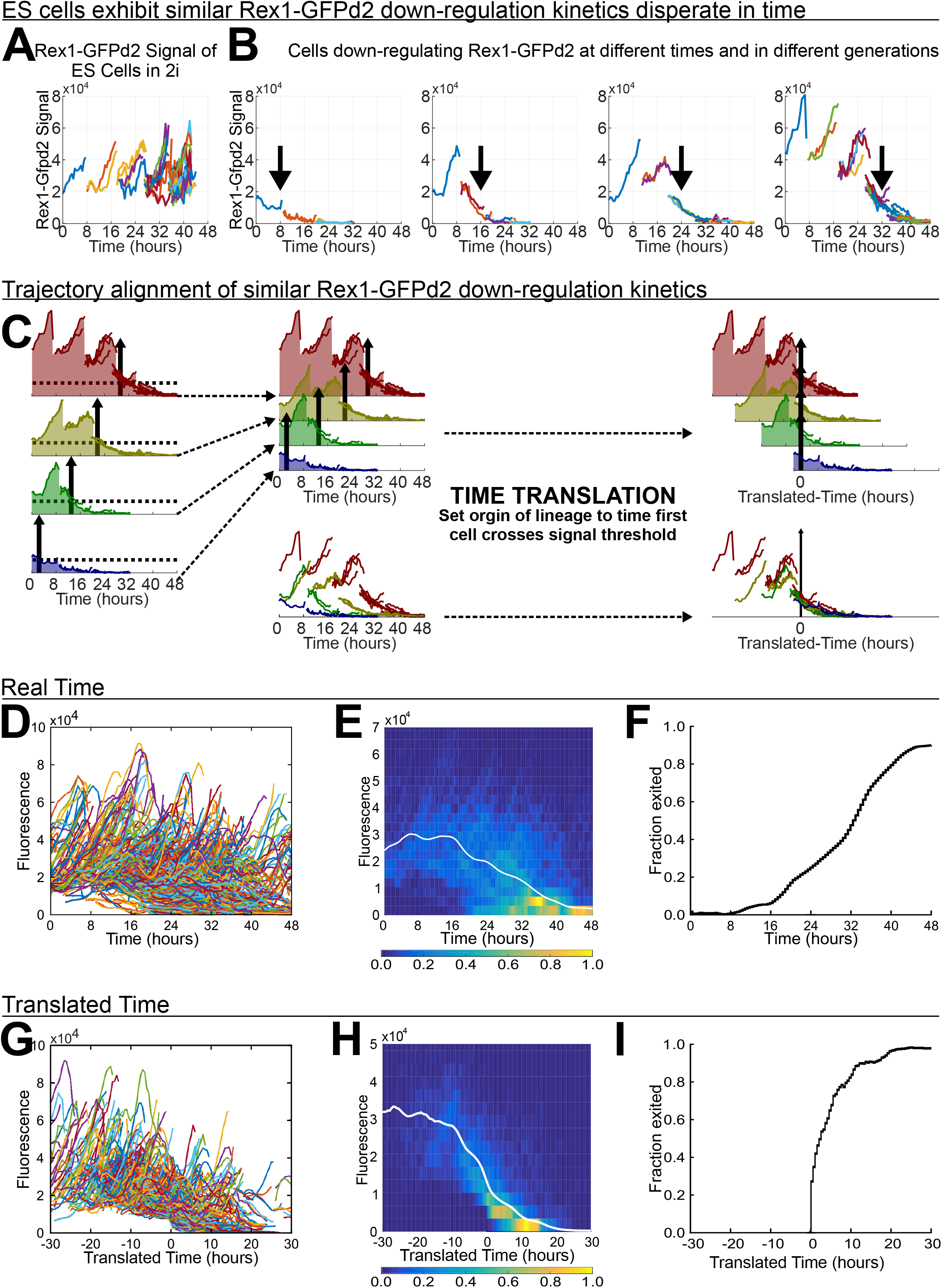
**A.** Rex1-GFPd2 trajectory for a representative ES cell lineage self-renewing in 2i. Each line is a cell. **B.** Rex1-GFPd2 trajectory for 4 representative lineages exiting in N2B27, downregulating the reporter at different times and in different generations. Black arrows indicate the Last generation of cells that are Rex1-GFPd2 positive. **C.** Schematic of the time-translation method for trajectory alignment. **D.** Real-time trajectories for 1,384 cells from 64 lineages in the N2B27 condition superimposed. **E.** Heat map of the real time trajectories for all lineages in the N2B27 condition with the time average shown as a white line. Colours indicate population density, ranging from 0 to 1. **F.** Real time percentage of cells that have crossed the signal threshold for all lineages. **G.** Translated time trajectories for all lineages in the N2B27 condition superimposed. **H.** Heat map of the translated time trajectories for all lineages in the N2B27 condition with the time average shown as a white line. Colours indicate population density, ranging from 0 to 1. **I.** Translated time percentage of cells that have crossed the signal threshold for all lineages.

To determine whether rapid decay in GFP was consistent across different lineages, we aligned all trajectories by a simple time-translation, which set the origin of time for each lineage to the moment the first cell of that lineage crossed the threshold for loss of GFP (black arrows in Fig. 6C). We compared the real time kinetics (Fig. 6D-F) to the translated time kinetics (Fig. 6G-I) by superimposing all tracked lineages, calculating the mean GFP trajectory for the population and then examining the exit kinetics on a per-cell basis. A reduction in GFP signal heterogeneity in time was observed when the time translation was performed (Fig. 6D,G), and the exponential decay phase became apparent at the level of the entire population (Fig. 6E,H). Time-translation was performed on entire lineages and yet this resulted in a strong reduction in variability of profiles for all individual cells analysed. All cells down-regulated GFP intensity by 50% within 3 hours of the first cells exiting, further indicating that cells in the same lineage exit naïve pluripotency in a synchronous fashion.

Finally, we compared the proportion of cells that have exited between real time and translated time and found the exit kinetics changed from a sigmoidal (Fig. 6F) to a hockey stick curve (Fig. 6I). Therefore, down-regulation of GFP occurs acutely in all cells. In other words, once cells initiate the process of exit, observed by silencing of Rex1-GFP expression, they all do so with fast and similar kinetics.

## DISCUSSION

In this study we applied continuous live cell imaging to examine the exit of mouse ES cells from naïve pluripotency in defined conditions, a system that recapitulates aspects of epiblast progression in the peri-implantation embryo (Kalkan et al., 2017; Smith, 2017). We tracked over 2500 cells in two replicate experiments. Analyses of these movies reveals that prior to exit, cells exhibit a modest shortening of the cell-cycle and a significantly larger cell surface area but with limited motility. Once cells have exited, the cell-cycle shortens further, cells contract but remain flattened, and motility increases. We did not observe any re-expression of the naïve state reporter after down-regulation, consistent with an irreversible exit process. In line with previous observations in mass culture, we saw that the exit of individual clones was asynchronous over a period of 48 hours. However, synchrony was observed between lineally related cells, indicating symmetry of fate choice. We showed that asynchronous differentiation via symmetric divisions is consistent with a simple population dynamics model. Asynchrony arises due to a variable initial lag phase of 15.04±10.56 (SD) hours preceding an acute transition that follows a rapid, exponential decay.

We identify physical parameters for tracking the ES cell transition process. Prior to transition, cells flatten and spread. Only after exit, however, is there a significant increase in cell motility. Generation times reveal that cell division rate also begins to shorten prior to exit from naïve pluripotency. Cell cycle shortening may therefore be an early marker of differentiation. as suggested in a recent report (Waisman et al., 2019). However, self-renewing ES cells in serum/LIF exhibit a faster cell cycle than 2i cells, with a shortened G1 phase (ter Huurne et al., 2017). It is possible therefore that the initial cell cycle change is a direct response to active Erk signalling and may be inconsequential for the differentiation process. Our data also show that cell cycle rate further shortens after exit, which may relate to the increased epiblast proliferation observed in the embryo after implantation (Snow, 1977).

In contrast to observations of ES cells in serum (Toyooka et al., 2008), we did not detect any instance of reactivation of the Rex1 reporter following down-regulation. Serum perturbs expression of the naïve transcription factor network (Chambers et al., 2007; Filipczyk et al., 2015; Hayashi et al., 2008; Toyooka et al., 2008), and we surmise that this disrupts the tight connection of Rex1 expression to naïve identity that obtains in the mouse embryo (Kalkan et al., 2017). In controlled culture conditions, however, down-regulation of Rex1-GFPd2 marks irreversible exit from naïve pluripotency.

Asymmetric fates are observed after cell division in many stem and progenitor cell systems (Simons and Clevers, 2011). Asymmetric ES cell fates have been observed in vitro in response to spatially localised Wnt signals (Habib et al., 2013). Previous live imaging studies have suggested that asymmetric divisions occur during differentiation of heterogeneous populations of ES cells (Nakamura et al., 2018). However, we did not observe asymmetric divisions in clonal density cultures initiated from 2i. On the contrary, we found that the Rex1-GFPd2 reporter followed very similar expression dynamics between sister cells, indicating that the predominant mode of exit is via symmetric divisions. This finding is supported by population dynamics modelling, which describes exit as a time-dependent process occurring via symmetric divisions. We also observed a high correlation between sister cell division times, consistent with previous studies (Cannon et al., 2015; Nakamura et al., 2018; Waisman et al., 2019).

Importantly, however, exit times between unrelated cells are uncorrelated, explaining the population asynchrony. Exit occurred at different times over the life of cells and within all generations analysed. Notably, however, time-translated alignment of lineages revealed a common sudden exponential decay in reporter expression. Thus, exit occurs rather abruptly but after a variable lag period. A focus for future studies will be to identify how this variability is determined. It is already known that prior exposure to LIF delays transition, likely by boosting expression of naïve transcription factors. One possibility, therefore, is that fluctuations in expression levels of naïve transcription factors in 2i may result in different starting points for network collapse. It also seems likely that cell to cell variability in Erk signalling, a major driver of transition, may contribute to different lag periods (Albeck et al., 2013; Aoki et al., 2013). Indeed, transition is accelerated by reducing negative feedback in the Erk pathway (Nett et al., 2018). Our mathematical model will also allow us to study whether other factors, such as the position of a cell in a colony or the local cell density, affect the lag periods.

Finally, we note that asynchronous transition is evident in the epiblast during the peri-implantation period in utero (Acampora et al., 2016; Malaguti et al., 2019; Neagu et al., 2020). Staggered exit may be a safeguard against premature exhaustion of naïve epiblast, which consists of only 10-12 cells in the mouse blastocyst, or be a mechanism for developing heterogeneity in the formative epiblast.

## METHODS

### Cell Lines

Rex1-GFPd2 ES cells carrying a destabilised green fluorescent protein (GFPd2) reporter in the Rex1 *(Zfp42)* locus have been described previously (Kalkan et al., 2017). The Gap43-mCherry reporter was created by introducing the palmitoylation sequence of Gap43 to the N-terminus of the mCherry sequence via PCR amplification using the primers Gap43_F: AAAAGTCGACTGCCACCACCATGCTGTGCTGTATGAGAAGAACCAAACAGGTTGAAAAGAATGATGAGGACCAAAAGATCATGGTGAGCAAGGGCGAGGAGGATA and mCherry_R: TTTTCTCGAGTTACTTGTACAGCTCGTCCATGCCG. The mCherry vector was a kind gift from Dr.

Paul Bertone. GAP43-mCherry was stably expressed in ES cells by co-transfection of pPB-CAG-Gap43-mCherry-pgk-hph with PB transposase (4:1 ratio) using Mirus TransIT LT1 lipofection reagent following the manufacturer’s protocols (MIR 2300). Transfected cells were selected using 200 μg/ml hygromycin in 2i + 10% FBS for a minimum of 7 days for stable PB integration. Individual clones were picked for brightness and expanded.

### Embryonic stem cell culture

Cells for all assays were maintained in humidified incubators kept at 37°C in 7.5% CO_2_ and 5% O_2_ in N2B27 medium supplemented with two inhibitors without LIF, as described in Mulas et al., 2019. Low O_2_ was used for improved cell viability during differentiation and recovery (Mazumdar et al., 2010). Cells were seeded onto plates coated with 0.1% (w/v) Gelatin (Sigma-Aldrich, G1890) in PBS for routine culture and plates coated with 5 ng/mL Laminin (Sigma-Aldrich, L2020) in PBS for all assays. Cells were passaged as single-cells using ESGRO Complete Accutase (Millipore, SF006).

### Factor Withdrawal and Recovery Assays

#### Clonal Recovery Kinetics

Cells were plated at a density of 500 cells/10 cm^2^ in 2i on laminin coated Falcon 6-well clear flat bottom multiwell plates (Corning 353046). After 12 hours, the medium was aspirated and replaced with N2B27. 2i was serially reapplied for the different time-points. A positive control of naïve cells in 2i was maintained throughout the withdrawal-time course. Cells were allowed to recover for at least 24 hours before being stained for DNA (1:1000 Hoechst33342 ThermoFisher, H3570). Plates were scanned for Hoechst and Rex1-GFPd2 using an Olympus IX51, Prior stage and Lumen light source, with CellSens software, using a 10x objective. Plates were then stained for alkaline phosphatase (Sigma, 86 R-1KT) and rescanned with transmitted light. Images were analysed by manual counting and scoring colonies using the ImageJ Plugin>Analyze>Cell Counter (Rasband et al., 1997).

#### Population-Level Recovery Kinetics

Cells were plated at a density of 5000 cells/10 cm^2^ in 2i on laminin coated Falcon 6-well plates. After 12 hours, the medium was aspirated and replaced with N2B27. 2i was serially reapplied for the different time-points. A positive control of naïve cells in 2i was maintained throughout the withdrawal-time course. Cells were allowed to recover for at least 24 hours before being harvested for analysis by flow cytometry. ES cells were dissociated to single-cells with accutase, then resusped in either 2i or serum/Lif+PD03. All samples were analysed with the CyAn ADP Analyser (Beckman Coulter), with 488nm laser coupled with fluorescein isothiocyanate (FITC) 530/40nm and phycoerythrin (PE) 575/25nm band pass filters. Flow profiles were analyzed and quantified with Summit V4.3 (Dako Colorado, Inc.).

#### Time-of-Exit Regression and Quantification

A 2-parameter weibull cumulative distribution function, *CDF*(*x*) = 1 – *exp*[(*x/λ*)^*γ*^], was fit to the time-course of the fraction of Rex1-GFPd2 negative colonies or AP positive colonies, for the clonal recovery kinetics, or the fraction of Rex1-GFPd2 positive cells, for the population-level recovery kinetics, using LSQCURVEFIT in MATLAB 2019a (MathWorks, Inc.). The mean and standard deviation for time-of-exit were calculated from the 2-parameter Weibull *CDF*. Empirical data and regression curves were plotted in Prism 7.00 (GraphPad Software, Inc.).

### Proliferation Kinetics

Cells were plated at a density of 500 cells/10 cm^2^ in 2i on laminin coated Falcon 6-well plates. After 12 hours, the medium was aspirated and replaced with N2B27. A positive control of naïve cells in 2i was maintained throughout the withdrawal-time course. At each time point cells were stained for DNA (1:1000 Hoechst33342). Plates were scanned for Hoechst using an Olympus IX51 Prior stage and Lumen light source, with CellSens software, using a 10x objective.

Once images were acquired, calibration curves were generated from fluorescent images by manually segmenting randomly selected nuclei. Linear regression was performed on the total intensity for the different number of nuclei to obtain the equation of the line. Calibration curves were generated for each plate by taking an equal number of nuclei from each well.To determine the number of nuclei per colony, the total fluorescence of the colony was acquired by manual segmentation in Fiji. The resulting value was background-corrected by the mean intensity of a local region without cells. This fluorescence value was then used to calculate the nuclei within that colony. Doubling time was then calculated from the log-linear slope of the growth curve (doubling time = log_o_(2)/slope).

### Long-term single-cell imaging

Cells were plated on laminin coated Hi-Q4 dishes (Ibidi 80416) at single-cell density (500 cells/ 10 cm2) in 2i and placed in humidified incubators maintained at 37°C in 7.5% CO_2_ and 5% O_2_. After 12 hours, the medium was aspirated and replaced with N2B27 for differentiation conditions or 2i for control conditions. The culture dish was returned to the 37°C humidified 7% CO_2_ and 5% O_2_ incubator for one hour to allow for the medium to equilibrate prior to imaging.

Time-lapse imaging was performed using a Leica SP5 confocal microscope with the Leica 20x 0.7NA dry lens (Leica, 506513). Seven adjacent 3.7 nm z-stacks were acquired at multiple positions for GFP (488nm excitation with Ar laser, 500-550nm emission detected by HyD2 at 150% gain and acousto-optic tunable filter (AOTF) at 15), mCherry (594nm excitation with HeNe laser, 600-630nm emission detected by HyD4 at 100% gain and AOTF at 12), and bright field at 30 minute intervals for 72 hours. Laser power was set at 30% and pinhole set at 2 Airy units. The microscope was enclosed in a temperature controlled chamber, ‘The Box, heated by ‘The Cube, (Life Imaging Sciences), and maintained at 37°C in humidified 7% CO_2_ and 5% O_2_.

### Semiautomated lineage tracking

Manual cell tracking and segmentation was performed using Tracer (England et al., 2006). Analyses of cell centroid tracks, cell shapes, Rex1-GFPd2 fluorescence quantification and genealogical relationships were performed using the tracks analysis software bundle oTracks (Blanchard et al., 2009), for which a new ES-cell specific module was written (see Supplemental Methods for code and further details) in IDL (Interactive Data Language, Harris Geospatial Solutions). (See Supplemental Methods for further details).

### Formal analysis

Formal analysis of the single-cell tracking data output from oTracks was performed using a custom CLASSDEF written in MATLAB 2019a (MathWorks, Inc.). This includes comparison of generational distributions, genealogical correlations, and multi-trajectory time-translations. (See Supplemental Methods for further details).

### Mathematical modelling and parameter inference

Model simulations and parameter inference were performed in MATLAB 2019a. Parameters were inferred by constrained optimisation using FMINCON over the sum squared error between the number of Rex1-GFPd2 positive and negative cells (see Supplemental Model for further details).

### Data availability

All code generated for this study, all the raw data and process genealogical information will be made available on GitHub.

## AUTHOR CONTRIBUTIONS

Conceptualisation SES, HK, GM, AS. Experimental design SES and GM. Investigation SES. Tracking analysis software development GBB and SES. Lineage tracking and Formal analysis SES. Mathematical Modelling SES and HK. Writing SES, HK, GM, AS. Supervision HK, GM, AS.

## ACKNOWLEDGEMENTS

Wellcome-MRC Cambridge Stem Cell Institute Core Facilities, especially support from Peter Humphreys with imaging and from Andy Riddell with flow cytometry. Richard Adams for adapting the Tracer tool used in the semi-automated lineage tracing. Dr. Tuzer Kalkan, Dr. Amy Li, Dr. Carla Mulas, Dr. Masaki Kinoshitam and Dr. Paul Bertone for discussions and technical assistance with experiments. Dr. Alexander G. Fletcher for technical assistance with the parameter inference and both Dr. Fletcher and Dr. Sara-Jane Dunn for helpful feedback on the mathematical modelling and the manuscript. Microsoft Research for PhD-studentship funding for SES. AS is a Medical Research Council Professor. G.M.’s laboratory is supported by grants from the Giovanni Armenise-Harvard Foundation, the Telethon Foundation (TCP13013) and an ERC Starting Grant (MetEpiStem). HK acknowledges support by the Horizon 2020 research and innovation programme for the Bio4Comp project under grant agreement number 732482 and by the Israel Science Foundation (grantNo. 190/19).

