## Supplemental Figures and Methods for "Embryonic stem cells commit to differentiation by symmetric divisions following a variable lag period"

**SUPPLEMENTAL MATERIALS**

**SUPPLEMENTAL FIGURES S1-S3**

**SUPPLEMENTAL TABLES 1 and 2**

**SUPPLEMENTAL METHODS**

**SUPPLEMENTAL MODEL**

**SUPPLEMENTAL REFERENCES**

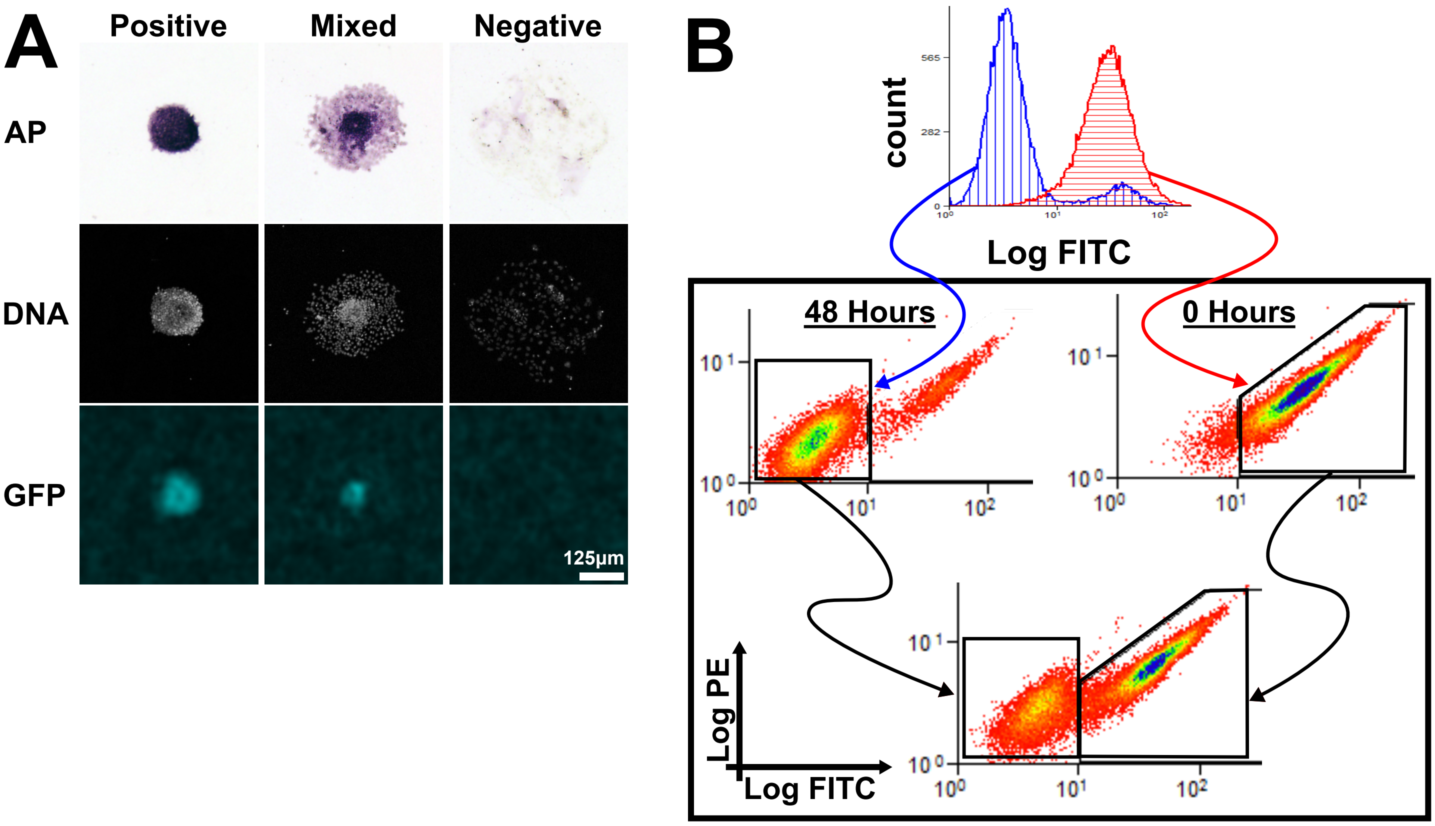
A
Positive
Mixed
Negative
AP
DNA
GFP
125µm
B
count
565
282
0
Log FITC
48 Hours
0 Hours
10 1
10 0
10⁰
10¹
10²
Log PE
Log FITC

**Figure S1 caption**

**A.** The clonal assay was analyzed by qualitative scoring of the colony by alkaline phosphatase staining (AP), top row; Rex1-GFP1d2 fluorescence, bottom row; and morphology. The middle row shows nuclei staining by Hoechst33342. The columns show representatives of the three types of colonies scored. A "positive" colony is compact and both AP and GFP positive, a "mixed" colony is larger with interspersed AP and GFP signal, and the "negative" colony is sparse with neither AP nor GFP expression. These representative images are cropped directly from Fig. 1C.

**B.** Flow cytometry gating for Rex1-GFPd2 positive and negative populations was performed using the positive control (red histogram) and final time-point (blue histogram), respectively. Black Box: Gates were defined on the two dimensional FITC v PE histogram. The Rex1-GFPd2 negative gate was made by construing a box around the population cloud in the lower-left corner of the 48 hour sample (top left). The Rex1-GFPd2 positive gate was made by constructing a non-overlapping polygon around the second distinct cloud above 1000 AU FITC (top right). These gates were then used for all other time-points within the same biological replicate (Example gating shown at the bottom).

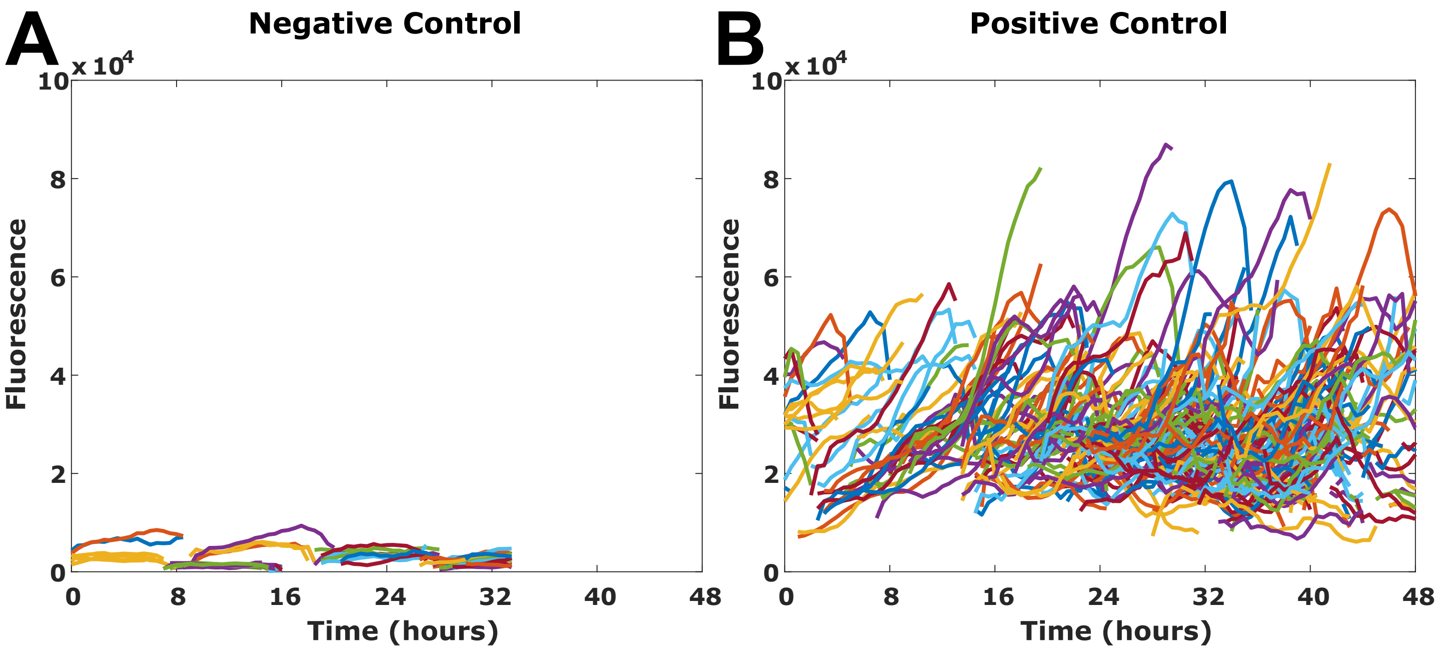


C

| Sample N | | Experiment 1 | | | Experiment 2 | | |
|---|---|---|---|---|---|---|---|
| Cell Line | Condition | Cells | Lineages | Data-Points | Cells | Lineages | Data-Points |
| Rex1-GFPd2 | N2B27 | 1,384 | 64 | 19,138 | 573 | 43 | 8,691 |
| Rex1-GFPd2 | 2i | 405 | 17 | 6,141 | 154 | 18 | 2,438 |
| E14 IVC | 2i | 56 | 6 | 664 | 52 | 6 | 1,258 |

**Figure S2 caption**

Cellular GFPd2 fluorescence trajectories for **A.** the negative control,E14 Tg2a ES cells in 2i, and **B.** the positive control, Rex1-GFPd2 ES cells in 2i. Each line represents an individual cell.

**C.** Summary of sample sizes for conditions and experiments.

### Doubling Time: 2i v N2B27

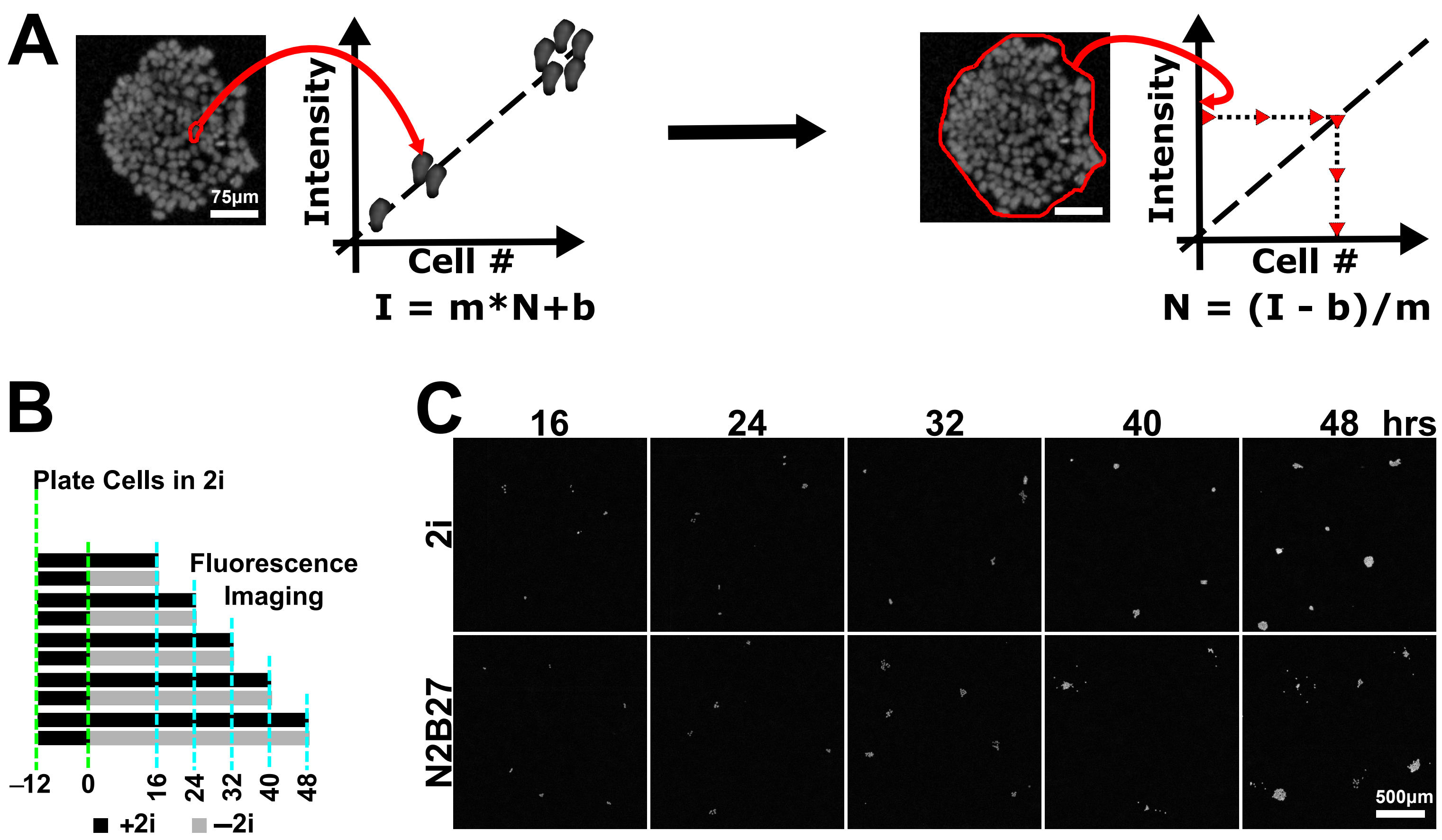


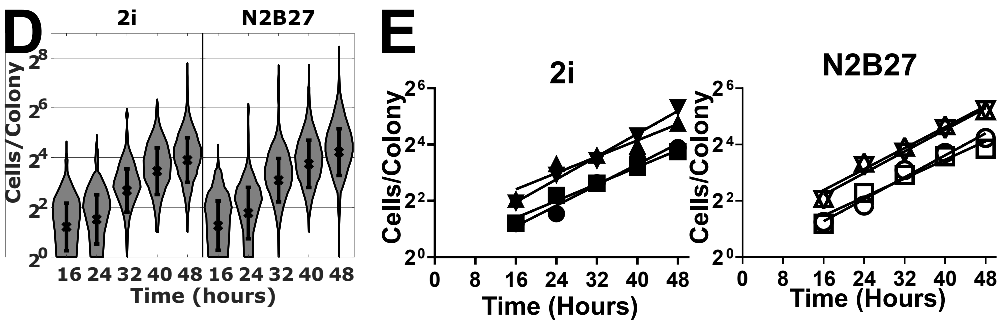


F

| Doubling Time (hours) | | ns |
|---|---|---|
| | 2i | N2B27 |
| Experiment 1 | 10.90 | 10.24 |
| Experiment 2 | 13.31 | 12.08 |
| Experiment 3 | 13.80 | 10.71 |
| Experiment 4 | 9.88 | 10.37 |
| Mean ± SD | 11.97±1.89 | 10.85±0.84 |

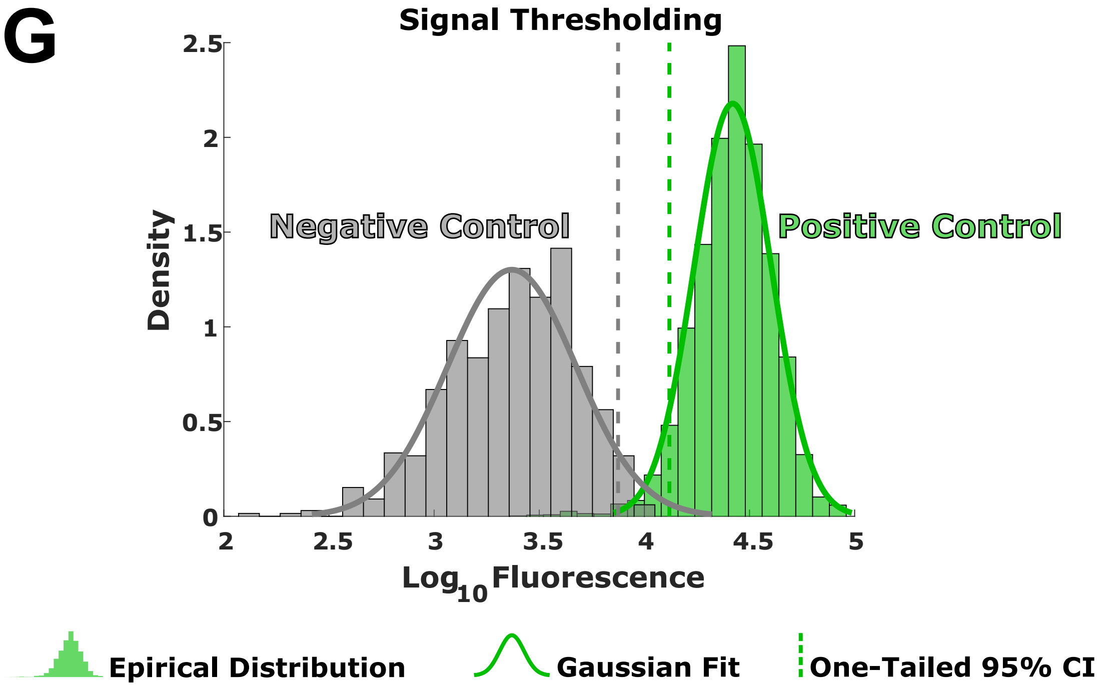

**Figure S3 caption**

**A.** Fluorescence based method for doubling-time quantification. The number of nuclei ***N*** per colony is determined with a linear calibration curve, parameterized by slope *m* and y-intercept *b*, for fluorescence intensity ***I***.

**B.** Experimental strategy used to measure the doubling time of cells cultured in 2i and N2B27.

**C.** Representative images from 1 out of 4 independent experiments of nuclei staining by Hoechst33342 after the indicated hours in 2i (top row) and N2B27 (bottom row).

**D.** Violin plots for the number of cells per colony for cells cultured in 2i (left) and N2B27 (right). The "x" shows the mean with error bars showing the standard deviation. One representative experiment out of 4 is shown.

**E.** Growth curves (solid lines) calculated from the mean number of cells per colony from each time point. Markers indicate experiment (n = 4).

**F.** The doubling time for cells cultured in N2B27 is numerically shorter than cells cultured in 2i, but not significantly (p-value = 0.3194, two-tailed unpaired t-Test) (n = 4).

**G.** Signal thresholding for Rex1-GFPd2 fluorescence was performed using the lower one-tailed 95% confidence interval of all segmented positive control cells (green, Rex1-GFPd2 ES cells in 2i). The distribution of the negative control (gray, E14 Tg2a ES cells in 2i) is shown as a reference.

### SUPPLEMENTAL TABLES

#### Supplemental Table 1

*Quantification of Time-of-Exit using 2-parameter Weibull CDF.* Parameters are reported ast the mean±SD (N=8).

| Marker | Mean (hrs) | SD (hrs) | $R^2$ | Scale (λ) | Shape (γ) |
|---|---|---|---|---|---|
| Rex1-GFPd2 (Population) | 31.5±3.5 | 13.2 ± 2.9 | 0.9701 ± 0.0155 | 35.4 ± 3.8 | 2.6 ± 0.6 |
| Rex1-GFPd2 (Clonal) | 32.1±3.4 | 20.0 ± 0.7 | 0.9660 ± 0.0132 | 35.9 ± 4.1 | 1.6 ± 0.2 |
| AP | 32.0±2.7 | 20.0 ± 1.7 | 0.9608 ± 0.0296 | 35.7 ± 3.2 | 1.6 ± 0.2 |
| Model | 29.6 | 9.7 | N/A | 33.0 | 3.37 |

#### Supplemental Table 2

*Comparison of Quantification of Time-of-Exit using 2-parameter Weibull CDF.* Adjusted P-values (Bonferroni correction) (N=8).

| Medium | Comparison | | Scale (lambda) | Shape (gamma) | Mean | SD |
|---|---|---|---|---|---|---|
| 2i | Population | GFP Clonal | 1 | 0.063 | 1 | 0.016 |
| 2i | Population | AP Clonal | 1 | 0.061 | 1 | 0.026 |
| 2i | GFP Clonal | AP Clonal | 1 | 1 | 1 | 1 |

### SUPPLEMENTAL METHODS

#### *Semiautomated lineage tracking*

The confocal z-stacks of the GFP channel, generated from long-term single-cell imaging, were summed into a single image per time point. To assist with manual segmentation, optimal slices of the  mCherry and bright field channels were manually selected. These three time series were concatenated into a 3-channel (Bright field, Gap43-mCherry, Rex1-GFPd2) 8-bit hyperstack (XYTC) and  imported into  Tracer (England et al., 2006) for manual cell tracking and segmentation. .

Lineage tracking and manual segmentation of cells  were performed on the 8-bit hyperstacks using the Tracer tool. Cells were identified by visual inspection of the different channels. Segmentation was performed free-hand by outlining the image of the tracking reporter with the mouse cursor. Once the outline had been completed, a digital mask was automatically generated over the outlined cell image, indicating that it had been tracked for that frame. Thus preventing double counting of both cells and pixels. Cells were also prescribed one of three fates: 'divided', 'died', or 'unknown'. The 'unknown' cell fate resulted when the cell moved out of frame or could not be reliably identified because of crowding. An additional mask which did not include any cells, named 'bkg', was generated for each image and the mean value of this mask was used to subtract the background from the corresponding image.

The output of the manual tracks from Tracer were used as the input for the analysis software bundle oTracks, with the 'CellFamilyFluorescenceIntensityQuantifier' function (Blanchard et al., 2009). This module performs analyses of cell centroid tracks, cell shapes, Rex1-GFPd2 fluorescence quantification and genealogical relationships. The output of oTracks is text files that pairs cell identity, cell position, 2d cell areas, fluorescence intensity, and genealogy for each image frame. Tracking and analysis software available upon request. The software was run on OS X Mavericks (Apple Inc.), using the VMware Workstation 10.0.1 (VMware, Inc.), hosted on Windows 8.1 Pro (Microsoft).

***Lineage reconstruction and parameter extraction***

Formal analysis of the single-cell tracking data output from oTracks was performed using a custom CLASSDEF in MATLAB 2019a (MathWorks, Inc.). The tracking data was compiled into a structure for biological cells named ‘cells’ and a structure of lineages time-courses called ‘lineages’. Each cell was paired with metadata to track the recorded values back to the cell in a video. These were: experiment date, 'date' = (yyyymmdd); well id, ‘well’ = (‘PosC', 'NegC', ‘Rep1’, ‘Rep2’); media,  ‘condition’ = (‘twoi’, ‘n2b27’); video number, 'position' = [1,30]; and lineage founding cell, ‘founder’ = [founder cell name]. All cells from unique lineages can be identified with these values.  Genealogical information was stored by the same convention as Blanchard et al., 2009. In brief, parent cells of a lineage were the cells observed in the first time frame of a video. Each cell in the field of view was given an arbitrary number starting from 1, e.g. ‘c1’ for the first recorded cell in frame one of a video. Daughter cells are named by appending the mother’s name with either an ‘a’ or a ‘b’ in an arbitrary manner, e.g. the daughters of ‘c1’ would be ‘c1a’ and ‘c1b’. From the naming, it is immediately possible to determine generation number and search for genealogical relationships. Further cell attributes from the oTracks or derived from this data are listed below:

**Supplemental Methods Table 1** Cellular attributes.

| Attribute | abbreviation | Calculation | units | Type |
|---|---|---|---|---|
| Fate | fate | | | string |
| Generation | generation | length(name) - 2 | | integer |
| Frame number of observation | frame | | | integer array |
| Time of observation | time | 0.5 hours × (frame - 1) | hours | float array |
| (x, y)-centroid | [x, y] | | µm | float array |
| Area | area | | $\mu m^2$ | float array |
| Mean Fluorescence | f | | abundance/$\mu m^2$ | float array |
| Total fluorescence | f_total | f × area | abundance | float array |

***Signal Thresholding***

Signal thresholding for Rex1-GFPd2 fluorescence was determined using the positive control, Rex1-GFPd2 ES cells in 2i. A lower one-tailed 95% confidence interval ($\alpha$=0.05) was generated from the log10 transformed total cellular fluorescence of all data-points. The mean, standard deviation and t-score were used to parametrically calculate the confidence interval. This value was then used as the signal threshold.

***Bulk and Generational Distribution Analysis***

Two-tailed unpaired Wilcoxon Rank Sum was used to compare medians for cell-cycle duration, average area, and average velocity between cells in 2i and N2B27 and between Rex1-GFPd2 positive and Rex1-GFPd2 negative cells in N2B27. For each parameter, we further specified the constraint that a cell would only be considered if both the birth and cell division was observed. Parameter calculations and constraint definitions are listed in

Supplemental Methods Table 2. Temporal trends, within each of these four groups, were assessed by grouping cells by generation. The Kruskal-Wallis Test, with Bonferroni post-hoc correction, was used for multiway comparison across generations within a condition. Generations that had fewer than three representative lineages were discarded from the analysis.

**Supplemental Methods Table 2** Bulk and generational distribution parameter calculations.

| **Parameters** | **Abbr.** | **Calculation** | **Birth Observed**<br><br>**generation > 0** | **Division Observed**<br><br>**fate == divided** |
|---|---|---|---|---|
| Cell-cycle Duration (hours) | life | time(end) - time(1) | true | true |
| Average Area ($\mu m^2$) | area_ba r | mean(area) | true | trrue |
| Average Velocity (µm/hour) | v_bar | $\frac{1}{N}\sum_{i=1}^{N-1}\sqrt{([x,y]_{i+1}-[x,y]_i)}$ | true | true |

***Genealogical Correlation Analysis***

The cell naming convention system is a binary string of 'a's" and "b's" (Blanchard et al., 2009; England et al., 2006) ( see Lineage reconstruction and parameter extraction in Supplemental Methods) was used to generate the genealogical comparisons **.** We employed an ancestor-to-descendant convention, for intergeneration comparisons, and an a-to-b convention, for intragenerational comparisons, i.e.. mothers-daughter {(c1 → c1a), (c1 → c1b)}, grandmother-granddaughter {(c1 → c1aa), (c1 → c1ab),(c1 → c1ba), (c1 → c1bb)}, sister-sister (c1a → c1b) , cousin-cousin {(c1aa → c1ba),(c1aa → c1bb),(c1ab → c1ba),(c1ab → c1bb)}. This unidirectional approach prevents double counting comparisons.

For each set of relationship-parameter set, we calculated a two-tailed paired Wilcoxon Rank Sum and Pearson's correlation coefficient. Significance for Pearson's correlation coefficient was determined via permutation test with replacement. With,

$$\text{p-value} = \frac{\text{(number of values more extreme than observation)} + 1}{\text{(number of permutations)} + 1} \ .$$

Parameter calculations for Rex1-GFPd2 signal used in the genealogical analysis are listed in Supplemental Methods Table 3, with the addition of the cell-cycle duration shown in Supplemental Methods Table 2.

**Supplemental Methods Table 3** Genealogical correlation parameter calculations

| **Parameters** | **Abbr.** | **Calculation** | **Birth Observed**<br><br>**generation > 0** | **Division Observed**<br><br>**fate == divided** |
|---|---|---|---|---|
| Initial Fluorescence | f_initial | f_total(1) | true | |
| Final Fluorescence | f_final | f_total(end) | | true |
| Average Fluorescence | f_bar | mean(f_total) | true | true |
| Integrated Fluorescence | f_AUC | trapz(time,f_total) | true | true |

**Time-translation Multi-trajectory alignment**

For the $i^{th}$ lineage, the time at which the first cell crossed the Rex1-GFPd2 signal threshold $t = \tau_i$ was determined. The entire time-course of the $i^{th}$ lineage was then translated such that time T($\tau_i$) = 0. All lineages were superimposed and summary statistics were generated from this data. Trajectory alignment was performed on a subset of lineages in which all cells were observed to have exited.

### SUPPLEMENTAL MODEL

#### Model derivation

Let $P$ be the population of Rex1-GFPd2 positive cells and $N$ be the population of Rex1-GFPd2 negative cells. Both cell types divide with a rate $\lambda$ and die at a rate that grows logistically with total cell number (specified by two parameters, $\delta$ and $K$). Additionally, $P$ cells have the ability to generate $N$ cells (i.e. exit naive pluripotency). We assert that exit is an irreversible process and we couple the exit process to the divisions of $P$. The divisions of $P$ are partitioned into symmetric self-renewal divisions with probability $\chi_R$, symmetric exit divisions with probability $\chi_S$, and asymmetric exit divisions with probability $\chi_A$. Where symmetric self-renewal divisions produce two ES cells, symmetric exit divisions produce two cells who have exited, and asymmetric divisions generate one cell of each type. These are the only division types allowed thus the sum of their probabilities is 1.

$$\chi_R(P,N;t) + \chi_S(P,N;t) + \chi_A(P,N;t) = 1. \quad (1)$$

This can be rewritten such that the divisions are partitioned into those that strictly generate more $P$ cells and divisions that give rise to at least one $N$ cell,

$$\chi_R(P,N;t) = 1 - f(P,N;t), \quad (2)$$

$$f(P,N;t) = \chi_S(P,N;t) + \chi_A(P,N;t). \quad (3)$$

We assume the probability of symmetric and asymmetric divisions are constant in time, which allows us to rewrite the probability of divisions that generate at least one $N$ as,

$$f(P,N;t) = \rho f(P,N;t) + (1-\rho) f(P,N;t), \quad (4)$$

where $\rho \in [0,1]$. We can now write the probability of each type of exit divisions in terms of $f$ as,

$$\chi_S(P, N; t) = \rho f(P, N; t), \quad (5)$$

$$\chi_A(P, N; t) = (1-\rho) f(P, N; t). \quad (6)$$

By observation, we have modelled the total cell number $C = P + N$ by logistic growth,

$$dC/dt = (\lambda - \delta \cdot (1 + \tfrac{C}{K_0})) \cdot C, \quad (7)$$

$$dC/dt = (\lambda - \delta - \delta \cdot \tfrac{C}{K_0}) \cdot C, \quad (8)$$

$$dC/dt = (\lambda - \delta)(1 - \tfrac{\delta}{(\lambda - \delta)K_0} \cdot C) \cdot C. \quad (9)$$

Let the net growth rate be $\gamma = \lambda - \delta$ and the carrying capacity be $K = \frac{(\lambda - \delta)K_0}{\delta}$, where $K_0$ determines how strongly cell death depends on cell density. Substituting these two terms in to (9) yields the growth term,

$$dC/dt = \gamma(1 - \tfrac{C}{K}) \cdot C. \quad (10)$$

The state transitions of the system are described as follows:

| **Dynamics** | $\{P, N\} \rightarrow$ | **Propensity** |
|---|---|---|
| Symmetric Renewal | $\{P+1, N\}$ | $(1-f) \cdot \lambda P$ |
| Symmetric Differentiation | $\{P-1, N+2\}$ | $\rho f \cdot \lambda P$ |
| Asymmetric Division | $\{P, N+1\}$ | $(1-\rho) f \cdot \lambda P$ |
| Negative Division | $\{P, N+1\}$ | $\lambda N$ |
| Positive Death | $\{P-1, N\}$ | $\delta(1 + \frac{N+P}{K_0}) \cdot P$ |
| Negative Death | $\{P, N-1\}$ | $\delta(1 + \frac{N+P}{K_0}) \cdot N$ |

Using these dynamics, we derive a coupled system of nonlinear ordinary differential equations which describe the time evolution of the two populations under exit conditions. First we derive the equation for the time derivative of positives cells $P$,

$$dP/dt \quad = \quad (1-f)\cdot\lambda P - \delta(1+\tfrac{N+P}{K_0})\cdot P - \rho f\cdot\lambda P\ , \tag{11}$$

$$= \quad [\lambda - \delta(1+\tfrac{N+P}{K_0})\ - \lambda f\ -\ \rho\,\lambda f]\cdot P, \tag{12}$$

$$= \quad [\lambda - \delta(1+\tfrac{N+P}{K_0})\ - (1+\rho)f\cdot\lambda]\cdot P. \tag{13}$$

Likewise, for the number of negative cells $N$,

$$dN/dt \quad = \quad \lambda N - \delta(1+\tfrac{N+P}{K_0})\cdot N + 2\rho f\cdot\lambda P\ + (1-\rho)f\cdot\lambda P, \tag{14}$$

$$= \quad [\lambda - \delta(1+\tfrac{N+P}{K_0})]\cdot N\ +\ (1+\rho)f\cdot\lambda\cdot P. \tag{15}$$

Substituting the the carrying capacity $K = ((\lambda - \delta)\cdot K_0)/\delta$ and net growth rate $\gamma = \lambda - \delta$ into (13) and (15), we arrive at,

$$dP/dt \quad = \quad [\gamma\cdot(1-\tfrac{N+P}{K}) - (1+\rho)f\cdot\lambda]\cdot P, \tag{16}$$

$$dN/dt \quad = \quad \gamma\cdot(1-\tfrac{N+P}{K})\cdot N + (1+\rho)f\cdot\lambda\cdot P. \tag{17}$$

**Variables and parameters:**

$P$, Rex1-GFPd2 positive cells.

$N$, Rex1-GFPd2 negative cells.

$\lambda$, division rate ($h^{-1}$).

$\delta$, basal death rate ($h^{-1}$).

$\gamma$, $\lambda - \delta$, net growth rate ($h^{-1}$).

$\chi_R$, probability of symmetric self-renewal divisions.

$\chi_S$, probability of symmetric exit divisions.

$\chi_A$, probability of asymmetric divisions.

$f$, $\chi_S + \chi_A$, the probability of an N producing division

$\rho$, ratio of symmetric divisions to all N producing divisions.

$K_0$, Density dependence of cell death.

$K$, Carrying capacity.

$c$, constant.

$n$, exponent.

$k$, hill function constant.

**Parameter Inference Constraints**

| Parameter | Constraint | Initial Guess |
|---|---|---|
| $\lambda$ | [log(2)/12, log(2)/10] | log(2)/12 |
| $\delta$ | [0,1] | 0.001 |
| $\gamma$ | [log(2)/13, log(2)/10] | log(2)/12 |
| $\rho$ | [0,1] | 0.5 |
| $K$ | [10,20] | 10 |
| $c$ | [0,1] | 0.01 |
| $n$, | [1, 3] | 2 |
| $k$ | [0,100] | 10 |

**Parameter inference for alternative test functions**

In the absence of further data, to model the probability of exit, we choose from a set of phenomenological candidate functions $f(P, N, t)$.

| Time evolution of Exit Probability | f(t) | ρ (Symmetry Bias) | BIC |
|---|---|---|---|
| Constant | c | 0.4624 | 20.52 |
| Ramp | $\min[c \cdot (P+N), 1]$ | 0.434 | 11.50 |
| | $\min[c \cdot t, 1]$ | 0.3404 | 11.95 |
| ***Leading-Order Polynomial*** | $\min[c \cdot (P+N)^n, 1]$ | ***1*** | ***8.68*** |
| | $\min[c \cdot t^n, 1]$ | ***0.9999*** | ***9.28*** |
| Hill Function | $c \cdot (P+N)^n/[(P+N)^n + k^n]$ | 1 | 12.50 |
| | $c \cdot t^n/[t^n+k^n]$ | 1 | 13.00 |

**Derivation of the weibull distribution using the time explicit leading order polynomial as the exit probability with respect to time**

The sum of (16) and (17) are:

$$\frac{d[N+P]}{dt} = \gamma(P+N)\cdot\left(1-\frac{N+P}{K}\right). \tag{18}$$

Which is the familiar differential form of the logistic growth equation (i.e. $\frac{dx}{dt} = cx\cdot(1-\frac{x}{k})$ ).

With initial conditions $P(0) = 1,\ N(0) = 0$, we solve (18),

$$P+N = \frac{K}{1+(K-1)\,e^{-\gamma t}}. \tag{19}$$

We eliminate N from (16) by moving $P$ to the RHS of (19) and substitute into,

$$dP/dt = \left[\gamma\cdot\left(1-\frac{1}{1+(K-1)\,e^{-\gamma t}}\right)-(1+\rho)f\cdot\lambda\right]\cdot P. \tag{20}$$

Using the time-explicit leading order polynomial for our exit function $f(t) = c\,t^{n}$,

$$\begin{aligned} dP/dt &= \left[\gamma\cdot\left(1-\frac{1}{1+(K-1)\,e^{-\gamma t}}\right)-c(1+\rho)\cdot\lambda t^{n}\right]\cdot P, \\ &= \left[\frac{\gamma}{1+\frac{e^{\gamma t}}{(K-1)}}-c(1+\rho)\cdot\lambda t^{n}\right]\cdot P. \end{aligned} \tag{21}$$

Integrating (21),

$$\begin{aligned} \int dP/P &= \int\frac{\gamma}{1+\frac{e^{\gamma t}}{(K-1)}}dt-\int c(1+\rho)\cdot\lambda t^{n}dt, \\ logP &= \int\frac{\gamma}{1+\frac{e^{\gamma t}}{(K-1)}}dt-c(1+\rho)/(n+1)\cdot\lambda t^{n+1}+c_0 \end{aligned} \tag{22}$$

The first term of the RHS can be integrated by two substitutions. First, we substitute $u = \gamma t$ and $du = \gamma dt$,

$$\int\frac{\gamma}{1+\frac{e^{\gamma t}}{(K-1)}}\,dt = \int\frac{1}{1+\frac{e^{u}}{(K-1)}}du. \tag{23}$$

Then, we substitute $s = e^u$ and $ds = e^u\, du$ and integrate by partial fractions,

$$\int \frac{1}{1+\frac{e^u}{(K-1)}}\, du = \int \frac{1}{s\cdot(1+\frac{s}{(K-1)})}\, ds,$$

$$= log\, s - log\,(\frac{s}{(K-1)} + 1)\quad ds. \tag{24}$$

Substituting back into (24) gives,

$$\int \frac{\gamma}{1+\frac{e^{\gamma t}}{K-1}} dt = \gamma t - log\,(\frac{e^{\gamma t}+K-1}{K-1}). \tag{25}$$

Thus, the analytical solution of (20) is,

$$P = \frac{K}{e^{\gamma t}+K-1}\cdot e^{\gamma t}\cdot exp[-\frac{c(1+\rho)\lambda t^{n+1}}{n+1}]. \tag{26}$$

The first two terms on the RHS of (26) comprise the logistic growth term and the final term is $1-[Weibull\ CDF]$. From the final term we are able to extract the shape and scale parameters for the Weibull distribution from our inferred parameters.

$$\lambda_{weibull} = \sqrt[n+1]{\frac{c(1+\rho)\lambda}{n+1}},\ (scale\ parameter), \tag{27}$$

$$\gamma_{weibull} = n+1,\ \ (shape\ parameter). \tag{28}$$
